# Human eIF2A has a minimal role in translation initiation and in uORF-mediated translational control in HeLa cells

**DOI:** 10.1101/2024.11.20.624465

**Authors:** Mykola Roiuk, Marilena Neff, Aurelio A. Teleman

## Abstract

Initiation of translation on eukaryotic mRNAs requires a 40S ribosome loaded with an initiator tRNA in order to scan for, and to identify, an initiation codon. Under most conditions, the initiator tRNA is recruited to the ribosome as part of a ternary complex composed of initiator tRNA, eIF2 and GTP. Although this function of recruiting the initiator tRNA was originally ascribed to another factor, eIF2A, it was later disproven and shown to belong to eIF2. Nonetheless, eIF2A is still considered a translation initiation factor because it binds the ribosome and shows genetic interactions with other initiation factors such as eIF4E. The exact function of eIF2A during translation initiation, however, remains unclear. Here we systematically test in HeLa cells, using ribosome profiling and luciferase reporter assays, the role of eIF2A in translation initiation, including translation of upstream ORFs that are either initiated with a AUG or near-cognate codons. Since eIF2A is thought to take over the function of eIF2 when eIF2 is inhibited, we also test conditions where the integrated stress response is activated, thereby leading to eIF2 inactivation. In none of our assays, however, could we detect a role of eIF2A in translation initiation. It is possible that eIF2A plays a role in translation regulation in specific conditions that we have not tested here, or that it plays a role in a different aspect of RNA biology.

## INTRODUCTION

For cells and organisms to grow and develop properly, they require a machinery that translates mRNAs in a finely-tuned manner. Alterations in the amount or activity of components of the translational machinery lead to severe pathological conditions (Tahmasebi et al., 2018). Of all the steps of protein production, translation initiation is considered to be the most precisely regulated and rate-limiting (Palmiter, 1975). Translation initiation under normal physiological conditions is well studied and occurs through a cap-dependent mechanism (Hinnebusch and Lorsch, 2012) that relies on the recruitment of the 40S ribosomal subunit preloaded with a set of initiation factors to the 5’-end of mRNAs. Correct positioning of the 40S at the 5’-end enables scanning, i.e. movement of the 40S in a 5’ to 3’ direction in search of the first initiation codon (AUG) within an optimal context (Pestova and Kolupaeva, 2002). The ribosome uses the CAU anticodon of the initiator Methionine tRNA (tRNA_i_^Met^) to identify the AUG start codon (Kozak, 1991). In the most common initiation pathway, tRNA_i_^Met^ is delivered to the 40S via the ternary complex which consists of eIF2 bound to GTP and the tRNA_i_^Met^ itself. However, several alternative initiation factors have been reported to possess the ability to also interact and potentially deliver initiator tRNA: eIF2D, the DENR/MCTS1 complex, and eIF2A (reviewed in (Grove et al., 2024)). We focus here on eIF2A, since its function is still elusive.

eIF2A was initially considered to be a functional analogue of prokaryotic IF2 (Shafritz and Anderson, 1970), however later this role was reassigned to the above-mentioned heterotrimeric factor eIF2 (α,β,γ) (Levin et al., 1973). Unlike eIF2, eIF2A was proposed to recruit tRNA_i_^Met^ to the 40S in a GTP-independent but codon-dependent manner (Zoll et al., 2002), however this activity was later challenged (Dmitriev et al., 2010) by attributing it to eIF2D which co-purified with eIF2A in the initial study. Unlike eIF2, eIF2A is dispensable for organismal viability, as single knock-outs of *eIF2A* in model organisms such as *Saccharomyces cerevisiae* (Komar et al., 2005)*, C. elegans* (Kim et al., 2018) and *M. musculus* (Anderson et al., 2021; Golovko et al., 2016) show little or no phenotype. However, double knock-outs of *eIF2A* and *eIF5B* exhibit severe growth defects both in yeast and in *C*.*elegans* (Kim et al., 2018; Zoll et al., 2002) and eIF2A is synthetic lethal with eIF4E in yeast (Komar et al., 2005). These genetic interactions suggest a role for eIF2A in translation initiation, however, the precise function of eIF2A is still not well defined, with contradictory results being reported. For example, eIF2A has been studied in the context of internal ribosome entry sites (IRES), where it was proposed to act both as a suppressor and an activator of IRES-mediated initiation. In yeast, eIF2A was reported to inhibit translation of URE2, GIC1 and PAB1 IRESes (Reineke et al., 2008; Reineke and Merrick, 2009). In Huh7 cells, knock-down of eIF2A was reported to hamper translation of a Hepatitis C virus (HCV) IRES reporter upon ER-stress (Kim et al., 2011), while a subsequent study detected no effect on the HCV-IRES reporter (Jaafar et al., 2016). Similarly, knock-out of eIF2A in HAP1 cells did not affect translation of either a HCV-IRES reporter or a EMCV-IRES reporter (Gonzalez-Almela et al., 2018), nor did it affect the infection rate of Sindbis virus (Sanz et al., 2019). Beyond IRES-mediated translation, eIF2A was reported to promote initiation from near-cognate start sites, initiated at leucine codons (CUG, GUG), by delivering elongator leucine tRNA^Leu^ to the 40S subunit (Sendoel et al., 2017; Starck et al., 2012; Starck et al., 2016). Such eIF2A-driven non-AUG initiation events were proposed to play a crucial role in different aspects of cell physiology and disease progression: cellular adaptation during the integrated stress response (Chen et al., 2019; Starck et al., 2016), fine-tuning of mitochondrial function (Liang et al., 2014), tumour progression (Sendoel et al., 2017), and repeat-associated non-AUG translation in familial amyotrophic lateral sclerosis (Sonobe et al., 2018). Such non-AUG initiation by eIF2A implies that eIF2A should possess high affinity for leucine tRNAs, however, according to *in vitro* filter binding assays, eIF2A binds inefficiently to elongator tRNA^Leu^ compared to initiator tRNA^Met^ (Kim et al., 2018), although, as mentioned above, it is questionable whether eIF2A has any tRNA binding capacity at all. Furthermore, loss of *eIF2A* in several systems did not recapitulate these effects on non-AUG initiation in either non-stressed or stress conditions (caused either by amino acid depletion or sodium arsenate treatment) (Gaikwad et al., 2024; Ichihara et al., 2021). In particular, one study (Gaikwad et al., 2024) found that eIF2A has little or no role in translation initiation and uORF mediated translation in yeast. Since in some cases mRNA translation in human cells can differ from mRNA translation in yeast, whether eIF2A also has such a minimal role in human cells remains to be clarified.

Due to these various discrepancies, we decided to study the role of eIF2A on mRNA translation and cellular fitness in human cells. For this, we generated *eIF2A* knock-out HeLa cells and applied ribosome profiling to analyze changes in mRNA translation. Overall, we find little contribution of eIF2A to cellular translation both under normal and stressed conditions in HeLa cells.

## RESULTS

### eIF2A-KO HeLa cell lines have no proliferative defect or change in global translation rates

To study the impact of eIF2A on cellular translation, we used CRISPR/Cas9 to generate two independent eIF2A knockout (*eIF2A*^KO^) HeLa cell lines, in which different exons of *eIF2A* were targeted. We confirmed the absence of eIF2A by immunoblotting (Fig.1A) and by genotyping the clones (Suppl. Fig. 1A). eIF2A^KO^ lines have reduced levels of *eIF2A* mRNA, likely arising from nonsense-mediated decay, due to premature stop codons in the *eIF2A* mRNA (Suppl. Fig. 1B). The eIF2A^KO^ cells exhibit no defect in cellular proliferation rates (Fig.1 B), which agrees with eIF2A having a dispensable role in organismal viability (Anderson et al., 2021; Golovko et al., 2016; Kim et al., 2018; Komar et al., 2005). To test if the loss of *eIF2A* affects global translation rates, we performed an *O*-propargyl-puromycin (OPP)-incorporation assay, which labels newly synthesized peptides and allows subsequent detection with anti-puromycin antibody. OPP-incorporation, however, did not show any significant change in *eIF2A*^KO^ cells compared to wild-type isogenic controls (Fig. 1C-D). In line with the OPP-incorporation assay we did not observe changes in polysome profiles of *eIF2A*^KO^ cells compared to controls (Fig. 1E-F).

**Figure 1:**
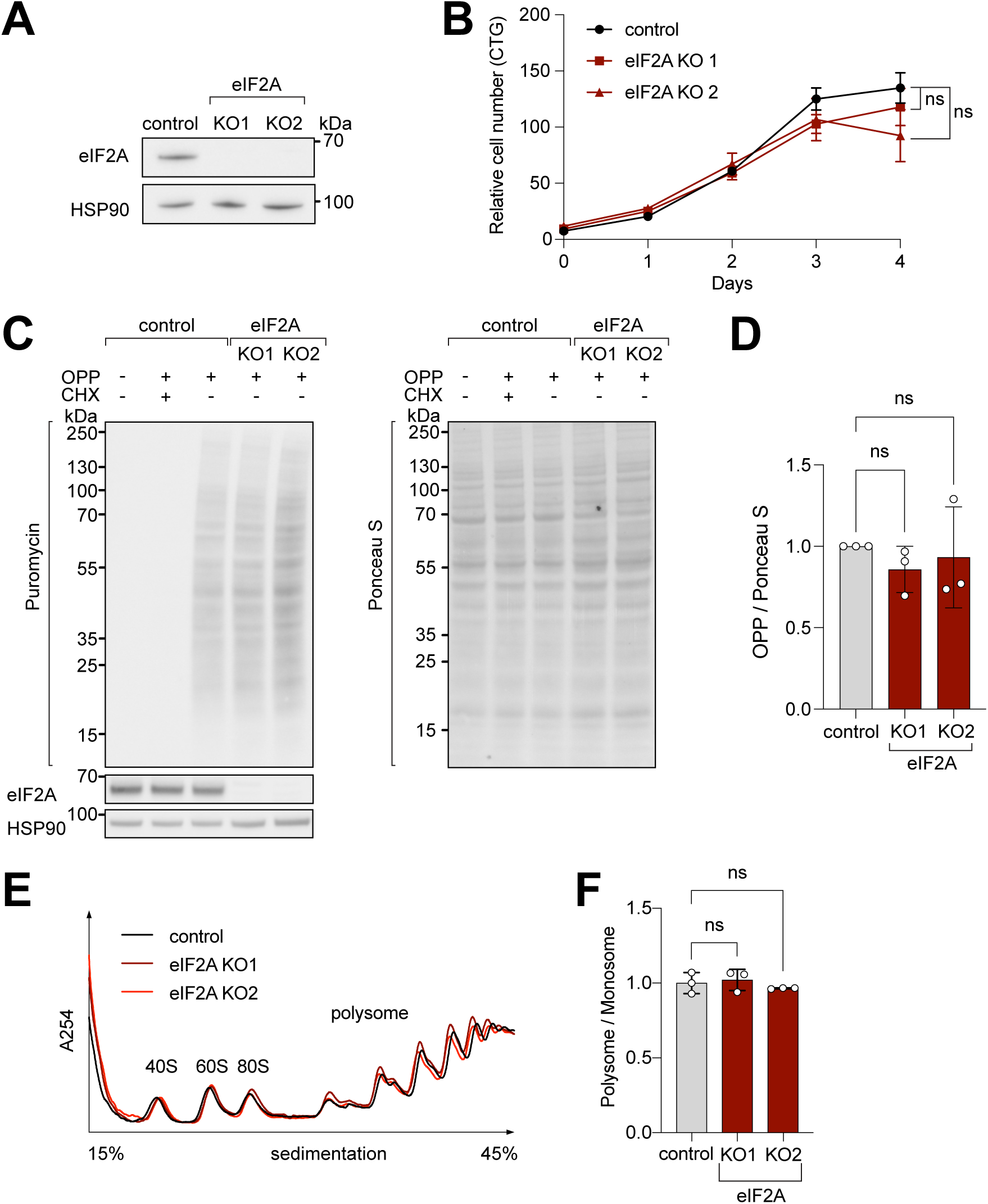
eIF2A has minimum effect on cell proliferation and global translation. **(A)** Validation that *eIF2A* knockout cells have no eIF2A protein by immunoblotting. **(B)** Two independent eIF2A knockout HeLa cell lines have no proliferation defect, assayed by CellTiter Glo. Error bars: standard deviation. Significance by ordinary one-way ANOVA. **(C-D)** eIF2A knockout HeLa cells have no detectable change in global translation rates compared to control cells. (C) The translation rate was measured by immunoblotting to detect OPP incorporated by metabolic labeling (left), and normalized to total protein amount assayed by PonceauS (right). Three independent replicates are quantified in panel (D). CHX = cycloheximide treated sample to completely block global translation (negative control). Error bars: standard deviation. Significance by ordinary one-way ANOVA. **(E-F)** Polysome profiles of *eIF2A*^KO^ cells show little to no difference to profiles from control cells. Lysates from either control or eIF2^KO^ HeLa cells were separated on a sucrose gradient. One representative graph is shown in panel D. The polysome/80S ratio of three independent replicates is shown in panel F. Error bars: standard deviation. Significance by ordinary one-way ANOVA.

Several studies have reported that stress can upregulate eIF2A levels (Panzhinskiy et al., 2021; Starck et al., 2016), or cause eIF2A to shuttle from the nucleus into the cytosol (Kim et al., 2011). Interestingly, Grove et al. recently reported (Grove et al., 2023) that increased levels on eIF2A in an *in vitro* translation system can suppress global translation initiation by directly binding and sequestering 40S ribosomal subunits. Thus, an increase in cytosolic eIF2A, either due to increased total levels or due to shuttling out of the nucleus, could be a potential mechanism to suppress translation upon stress. To determine if either localization or global levels of eIF2A change in response to stress in HeLa cells, we performed subcellular fractionation and loaded on a gel nuclear and cytosolic fractions in proportion to their abundance in a cell (i.e. both fractions were lysed in equal volumes). In HeLa cells eIF2A is present in both the cytosol and the nucleus, with higher levels in the cytosol (Suppl. Fig. 1C). Nonetheless, stress caused by tunicamycin treatment did not affect the subcellular distribution of eIF2A (Suppl. Fig. 1C-D), nor overall eIF2A levels (Suppl. Fig. 1F-G). To assess the localization of eIF2A with an orthogonal approach, we performed cell immunostaining. Since the eIF2A antibodies we have are not suitable for immunostaining, we overexpressed N-terminally FLAG-tagged eIF2A in HeLa cells treated cells with either DMSO or 100µM sodium arsenite (SA) for 1 hour (condition used in (Kim et al., 2011)). This revealed that overexpressed eIF2A was excluded from the nucleus, and showed a cytosolic, speckled staining pattern, which was not affected by SA treatment (Suppl. Fig. 1E). These results are consistent with what has been observed for endogenous eIF2A in HAP1 cells (Gonzalez-Almela et al., 2018; Sanz et al., 2017). To check if cellular stresses affects eIF2A levels in HeLa cells, we treated cells with different agents at concentrations and timeframes that were previously reported to alter eIF2A levels (Starck et al., 2016). However, we did not record significant changes in eIF2A levels with any of these treatments (Suppl. Fig. 1F-G). Nonetheless, we decided to test whether elevated levels of eIF2A would have the capacity to inhibit global translation levels *in vivo*. To this end we overexpressed eIF2A and quantified global translation rates by OPP incorporation but did not observe any significant change (Suppl. Fig. 1 I-H). In sum, our results indicate that loss of eIF2A as well as eIF2A ectopic overexpression in HeLa cells has little or no impact on global translation and cellular proliferation.

### Ribosome profiling identifies no eIF2A-dependent transcripts

We next investigated whether loss of *eIF2A* causes a change in translation of specific mRNAs which might be overlooked when assaying total global translation. To do this, we performed ribosome profiling, which sequences the mRNA footprints protected by ribosomes, on control and *eIF2A*^KO^ cells. Normalization of the footprint counts for each mRNA relative to the abundance of that mRNA in total RNA yields an estimate of a transcript’s translation efficiency. We used *eIF2A*^KO^ clone 1, since the exon targeted in this clone is further 5’ than in clone 2, thereby yielding a short, truncated protein of only 24 amino acids (Suppl. Fig 1A). Correlation analysis revealed that both footprint counts and total RNA counts were highly reproducible across technical triplicates in both control and *eIF2A*^KO^ cells (Suppl. Fig. 2A). In addition, our data revealed the expected enrichment of footprints in the coding sequence versus UTRs, as well as distinct triplet periodicity, both hallmarks of ribosome profiling data (Suppl. Fig. 2 B,C). Metagene analysis showed similar global profiles in footprint densities within ORFs in wild type and *eIF2A*^KO^ samples, in agreement with the data presented above that eIF2A is dispensable for global mRNA translation (Suppl. Fig. 2 B,C). A per-gene analysis identified only 15 genes in addition to eIF2A whose translation efficiency changes in *eIF2A*^KO^ cells compared to control cells (Fig. 2A). This is very different from our previous footprinting studies where we found that loss of DENR leads to reduced translation of 517 mRNAs (Bohlen et al., 2020), loss of PRRC2A/B/C leads to altered translation of 109 mRNAs (Bohlen et al., 2023), and expression of 4E-BP leads to altered translation of 605 mRNAs (Roiuk et al., 2024), suggesting that in comparison eIF2A plays a minor role in translational regulation (Fig. 2A). We decided to validate some of these eIF2A-dependent candidates by immunoblotting, and to our surprise, amongst all the proteins we tested, only CCND3 showed reduced levels in one of the two *eIF2A*^KO^ clones (Fig 2 B,C). We then quantified mRNA levels for all the genes that we tested (Fig. 2D). For NCAPH2, PPFIA1 and RPS6KB2, mRNA levels were either unchanged or reduced in the *eIF2A*^KO^ cells, indicating that translation efficiency for these genes was either unchanged or increased upon loss of eIF2A, which does not agree with the ribosome profiling data. For CCND3, protein levels were reduced in KO1 and mRNA levels were not, consistent with a drop in CCND3 translation in these cells, but this effect was not reproduced in eIF2A KO2 (Fig. 2C-D). Together, this indicates there is an enrichment for false-positives amongst the 15 genes identified by ribosome profiling. These results would be consistent with eIF2A having no effect on translation of any mRNA, with a few false-positives coming through in the transcriptome-wide ribosome footprinting, as is always the case. Nonetheless, to test further whether translation of any of these candidates is altered upon loss of eIF2A, we generated reporter constructs by cloning the 5’-UTRs of these candidates upstream of Renilla luciferase, and then testing their expression in the presence or absence of eIF2A. We used this approach multiple times in the past to identify mRNAs whose translation is dependent on initiation factors (Bohlen et al., 2023; Roiuk et al., 2024; Schleich et al., 2014). We succeeded in cloning 14 of the 15 5’ UTRs of the transcripts predicted to be eIF2A-dependent. None of the reporters, however, showed a consistent drop in translation in eIF2A KO1, eIF2A KO2, and siRNA-mediated eIF2A knockdown cells (Fig. 2E, Suppl. Fig. 3A-B). Although we did not exclude a possible effect of eIF2A on translation via the 3’UTR or coding sequence of the 12 genes that we did not assay via immunoblotting (Fig. 2B-D), our results indicate that eIF2A has little or no effect on translation of cellular mRNAs in HeLa cells under non-stressed conditions. Consistent with this, analysis of our ribosome footprinting data with anota2seq (Oertlin et al., 2019), also did not identify any transcripts with altered translation (Suppl. Fig. 3C).

**Figure 2:**
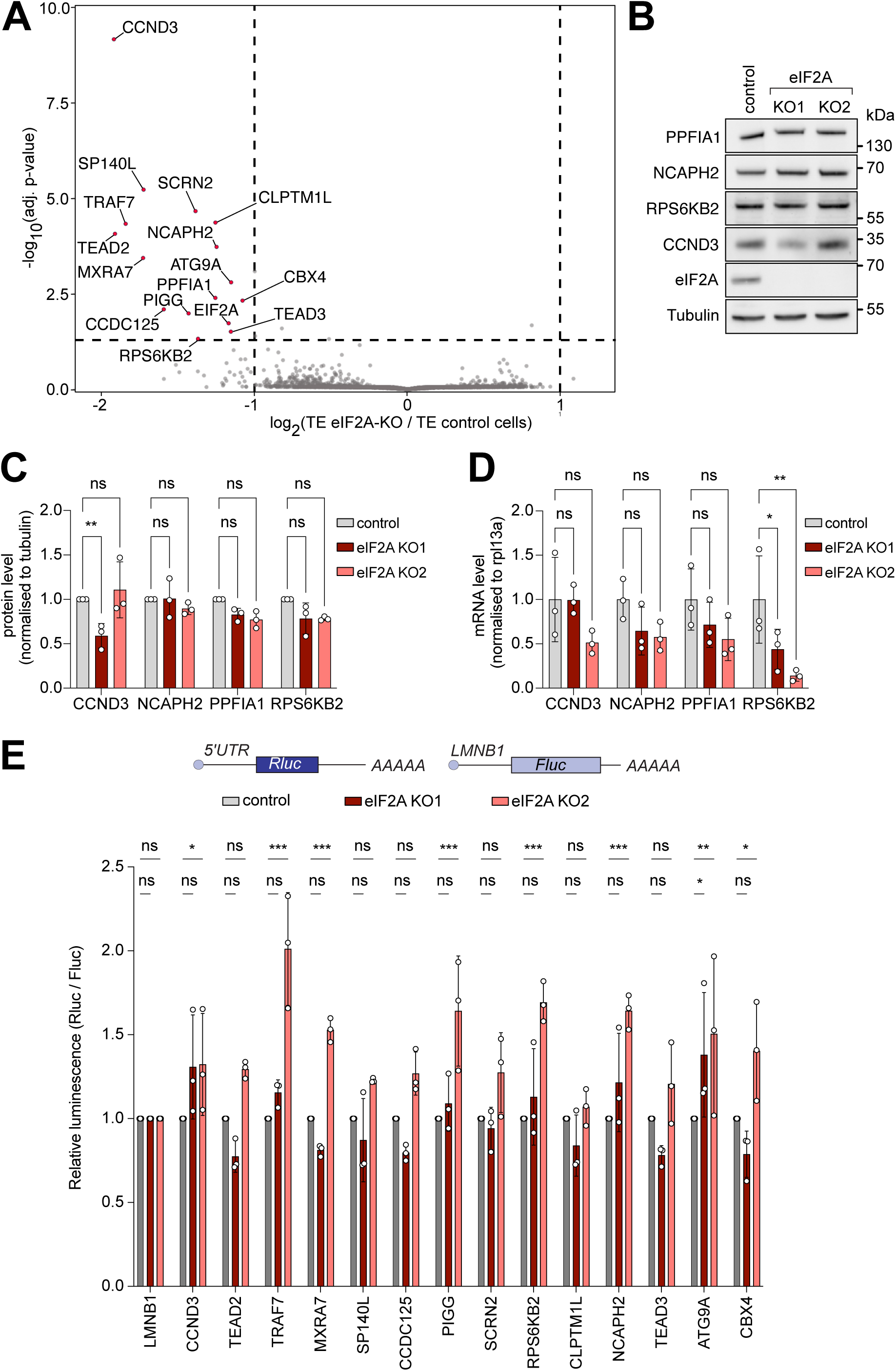
Ribosome profiling of eIF2A-KO lines finds little impact of eIF2A on translation. **(A)** Ribosome profiling identifies a handful of mRNAs sensitive to eIF2A depletion. Scatter plot of log2(fold change of Translation Efficiency eIF2A^KO^/ control) versus significance. Significant candidates with log2(fold change) < −1 are shown in red. Significance was estimated with the Wald test performed by the DESeq2 package. p-values are adjusted for multiple comparison. **(B-D)** Western blot validation of ribosome profiling results. Among the tested candidates, only CCND3 shows decreased protein levels in one eIF2A^KO^ clone. Representative blot in (B), of triplicates quantified in (C). mRNA levels of the corresponding transcripts are quantified and shown in (D). Significance by Dunnett’s multiple comparison test ANOVA. error bar = st. dev., ns = not significant, * p < 0.05, ** p < 0.01 **(E)** Luciferase reporters harboring 5’ UTRs of eIF2A-dependent transcripts do not show strong changes in expression upon loss of eIF2A. Reporters carrying the 5’ UTRs of the indicated candidate genes were cloned upstream of Renilla Luciferase (RLuc) and co-transfected with a Firefly Luciferase (FLuc) normalization control. The negative control RLuc reporter and the FLuc normalization control carry the 5’UTR of Lamin B1 (LMNB1). Significance by Dunnett’s multiple comparison test ANOVA. error bar = st. dev., ns = not significant, * p < 0.05, ** p < 0.01, ***<0.001

Since eIF2A was proposed to deliver tRNA to the ribosome in a codon dependent manner, we tested if loss of eIF2A affects translation initiation on a reporter where the ribosome is directly positioned on the initiation AUG. To perform this, we cloned a 5’ UTR reporter with a short 5’UTR (12 nt) where loading of the small ribosomal subunit should place the AUG in close proximity to the P-site (Gu et al., 2021). In addition, we tested a reporter bearing the EMCV IRES in the 5’-UTR, where initiation relies on close positioning of the 40S to the AUG (Davies and Kaufman, 1992). Translation of neither reporter, however, was affected by loss of eIF2A (Suppl. Fig. 3D). In sum, we were not able to find any transcript whose translation depends on eIF2A under non-stressed conditions in HeLa cells.

### eIF2A does not contribute to uORF translation

Several studies have reported that eIF2A can deliver alternative initiator tRNAs to uORFs with near-cognate start codons (Sendoel et al., 2017; Starck et al., 2012; Starck et al., 2016). Based on these reports we tested if eIF2A depletion affects uORF translation in HeLa cells. First, we analysed if translation initiation or termination on endogenous, AUG-initiated uORFs is altered in *eIF2A*^KO^ cells. For this, we calculated a metagene profile of footprint reads at start and stop codons of all AUG-initiated uORFs, however we could not detect any significant differences in uORF translation (Suppl. Fig. 4A,B).

Since uORFs are elements that inhibit translation of the downstream main ORF (mORF), initiation factors that influence uORF translation also influence, as a consequence, translation of the downstream main ORF. Therefore, we checked if translation efficiency of the main ORF of uORF-bearing transcripts changes between eIF2A^KO^ and control cells. For this we looked at the translation efficiency of different sets of transcripts: 1) all transcripts, 2) transcripts possessing AUG-initiated uORFs, or 3) transcripts containing uORFs starting with a near cognate initiation codon – CUG, GUG or UUG (Suppl. Fig. 4C). Although essentially no genes show a significant change in translation efficiency when analyzed singly (Fig. 2A), this aggregate analysis might identify significant trends caused by small changes in groups of transcripts. This analysis revealed, however, that translation of transcripts with near-cognate uORFs did not differ from the global distribution in translation efficiency of all transcripts (Suppl. Fig. 4C). Transcripts with AUG-initiated uORFs, however, did have a very slight, but statistically significant increase in translation efficiency, compared to the whole dataset (log2(fold change) of −0.03 versus −0.05) (Suppl. Fig. 4C).

To validate this minor role of eIF2A in uORF translation we tested various luciferase reporters in eIF2A^KO^ cells. We placed Renilla luciferase under control of a 5’ UTR with no uORF (LMNB1, as a negative control) or the same 5’UTR where we synthetically introduced a uORF with different start codons or Kozak sequences (Fig. 3A). In line with our ribosome profiling data, all of these luciferase reporters showed no significant difference in expression in *eIF2A*^KO^ or eIF2A overexpressing cells compared to controls (Fig. 3A, Suppl. Fig. 4D). Considering that this luciferase assay is a readout for translation of the main ORF, and not the uORF directly, we also performed an experiment where we directly visualised uORF peptide production. For this purpose, we generated fluorescent reporters carrying a uORF coding for the SINFEKL peptide which can be presented on the surface of cells expressing the H-2Kb Class I MHC complex (HEK293T-H2-Kb) where it can be detected with antibodies (Starck et al., 2016). To assess the impact of eIF2A on the production of this peptide, we generated a HEK293T-H2-Kb, *eIF2A*^KO^ cell line (Fig. 3B). Like *eIF2A*^KO^ HeLa cells, HEK293T-H2-Kb *eIF2A*^KO^ cells also did not display any defect in proliferation (Suppl. Fig. 4E) or global translation (Suppl. Fig. 4F-G). This revealed, however, no difference in translation of the uORF peptide when comparing *eIF2A*^KO^ to control cells, regardless of whether the uORF was initiated by a AUG codon or a near-cognate initiation codon, as detected by flow cytometry (Fig. 3D). This differs to what we previously observed upon loss of PRRC2A/B/C proteins, which caused increased expression of the uORF SINFEKL peptide and reduced expression of the downstream main ORF (Bohlen et al., 2023).

**Figure 3:**
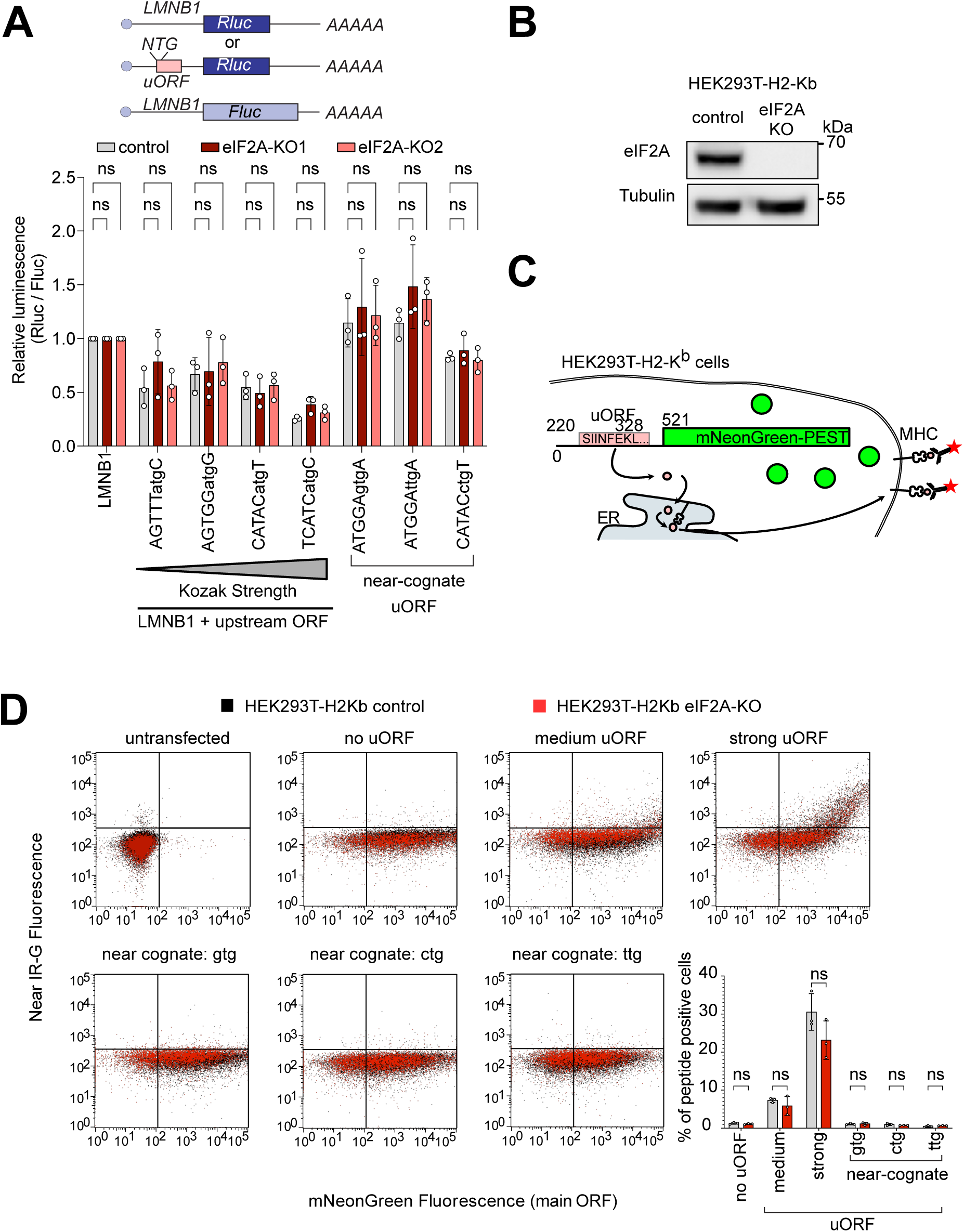
eIF2A has little or effect on uORF translation. (**A**) Synthetic reporters harboring uORFs with different start codons and initiation contexts do not show dependence on eIF2A. The sequence context of the uORF start codons is indicated: either AUG or a near-cognate start codon (GTG, TTG, CTG) was used. Significance by Dunnett’s multiple comparison test ANOVA, error bar = st. dev. ns = not significant. (**B**) Validation that HEK293T-H2-K^b^ *eIF2A^KO^* cells have no eIF2A protein by immunoblotting. (**C**) Schematic diagram illustrating the setup to simultaneously detect a small peptide produced by a uORF and fluorescent mNeonGreen encoded by the main ORF. The short peptide SIINFEKL is presented on the cell surface by MHC-I and detected using a monoclonal antibody. (**D**) *eIF2A* knockout does not cause a drop in uORF translation. In the graph to the right, the percent of uORF-positive cells relative to all mNeonGreen cells is quantified. Significance by unpaired, two-sided, t-test. ns = not significant.

### No detectable role for eIF2A in translation when eIF2 is inhibited

eIF2A has been proposed to act as a backup pathway for tRNA delivery when eIF2 function is attenuated due to phosphorylation of its alpha subunit by one of the kinases of the integrated stress response (Kim et al., 2018; Kwon et al., 2017; Starck et al., 2016; Tusi et al., 2021). We therefore induced the integrated stress response and tested the impact of loss of eIF2A on several different translational changes that occur – the global reduction in translation levels, the formation of stress granules, and induction of the few target genes such as ATF4 which evade the global reduction in translation and instead are translationally induced (Andreev et al., 2015; Sidrauski et al., 2015). For this we treated cells with tunicamycin or sodium arsenite to phosphorylate eIF2α (eIF2S1) by two independent kinases – PERK or HRI – and estimated global translation by assessing polysome profiles. Both treatments led to suppression of translation (compared Suppl. Fig. 5A-B to Fig. 1E) however the magnitude of suppression was similar in *eIF2A*^KO^ and isogenic control HeLa cells (Suppl. Fig. 5A-B). This suggests that, as in non-stressed conditions, eIF2A has a minimal effect on global translation also when the integrated stress response is active. We next tested if *eIF2A*^KO^ cells have any defect in stress granule formation, which is a hallmark of translation inhibition upon eIF2α phosphorylation (Sidrauski et al., 2015). However, eIF2A^KO^ cells formed stress granules to the same degree as the isogenic control HeLa cells (Suppl. Fig. 5C-D). To test induction of target genes during the integrated stress response we selected tunicamycin treatment because sodium arsenite is too strong and completely shuts off all translation. We cloned luciferase reporters carrying 5’UTRs of genes that were previously shown to be translationally upregulated upon eIF2α phosphorylation (Andreev et al., 2015). We then transfected these into control or *eIF2A*^KO^ HeLa cells and treated the cells with either DMSO or 1 µg/ml tunicamycin for 16 hours. Tunicamycin treatment resulted in the same level of eIF2α phosphorylation in either cell line (Suppl. Fig. 5E). None of the reporters we tested – *ATF4, PPP1R15B, IFRD1* – had an induction defect in *eIF2A*^KO^ cells (Fig. 4A). This is in line with a previous report showing that in yeast, the ATF4 homolog GCN4 is induced properly during amino acid starvation in cells lacking eIF2A (Zoll et al., 2002).

**Figure 4:**
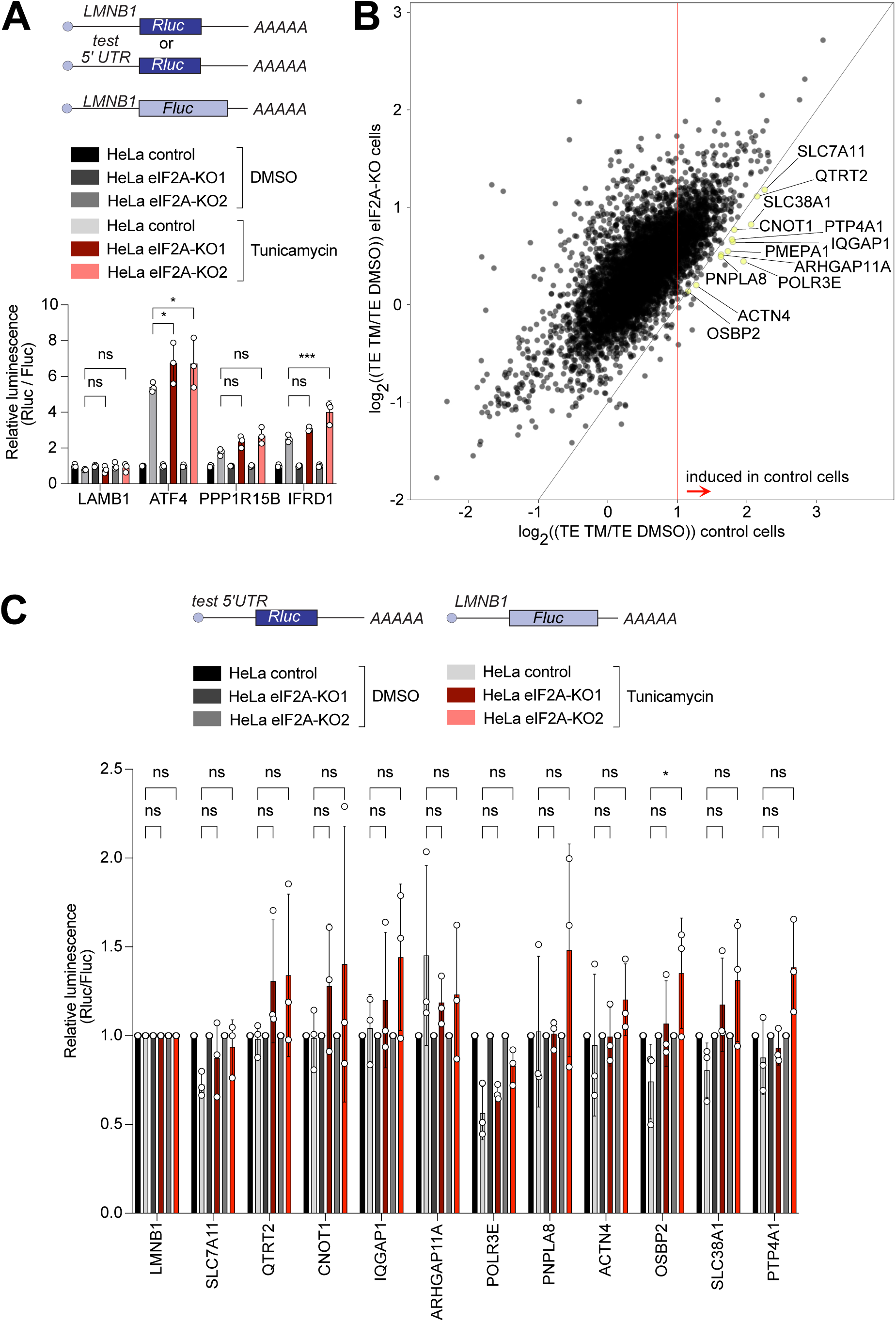
eIF2A has a minor impact on translation during the integrated stress response. (**A**) Loss of eIF2A does not blunt induction of target genes of the integrated stress response. Reporters carrying the 5’ UTRs of the indicated candidate genes were co-transfected with a Firefly Luciferase (FLuc) normalization control reporter into eIF2A^KO^ and control HeLa cells and treated for 16 hours either with DMSO or 1µg/ml tunicamycin (TM). Significance by Dunnett’s multiple comparison test ANOVA. error bar = st. dev., ns = not significant, * p < 0.05, ***<0.001 (**B**) Ribosome profiling identifies 12 mRNAs that are significantly induced upon tunicamycin treatment (1 ug/ml) in control cells but not eIF2A^KO^ cells. Scatter plot of log2(fold change) of Translation Efficiency TM/DMSO for control cells on the x-axis versus eIF2A^KO^ cells on the y-axis. mRNAs that are statistically significantly induced with log2(fold change)>1 in control cells but not in eIF2A^KO^ cells are shown in yellow and marked by gene name. Significance was estimated with the Wald test performed by DESeq2 package. p-values are adjusted for multiple comparison. (**C**) Transfection of luciferase reporters harboring 5’ UTRs of eIF2A-dependent transcripts do not show impaired induction in eIF2A^KO^ cells upon tunicamycin treatment. 5’ UTRs of eIF2A-dependent transcripts from panel B were cloned upstream of Renilla luciferase and co-transfected with a Firefly Luciferase (FLuc) normalization control reporter into control or EIF2A^KO^ HeLa cells with subsequent treatment for 16 hours either with DMSO or 1ug/ml tunicamycin (TM). Significance by Dunnett’s multiple comparison test ANOVA. error bar = st. dev., ns = not significant, * p < 0.05.

The assays described above mainly assess bulk translation, and might have missed changes in translation of individual mRNAs. To test whether eIF2A affects translation of any mRNA when eIF2 is inactive, we performed ribosome profiling on control and eIF2A^KO^ HeLa cells treated with tunicamycin and compared these data to the data from non-stressed controls described above. The results showed good reproducibility across duplicates for both ribosome profiling and total RNA samples (Suppl. Fig. 6A). Surprisingly, by comparing translation efficiency for all mRNAs in *eIF2A*^KO^ versus control cells in the tunicamycin-treated condition, we did not find any transcript with significantly changed translation apart from eIF2A itself (Suppl. Fig. 6B). In line with this, we also did not find any significant changes in a metagene profile of footprints on all main ORFs in control and *eIF2A*^KO^ cells upon tunicamycin treatment (Suppl. Fig. 6 C-D).

Nonetheless, it is possible that the induction of particular transcripts in response to stress may be affected in eIF2A-knock-out cells compared to controls. To test it, we compared changes in translation efficiency between untreated and treated cells in both control and *eIF2A*^KO^ cells. This identified a small group of transcripts whose translation was more strongly induced in control cells than in *eIF2A*^KO^ cells (Fig. 4B). Statistical analysis identified 12 transcripts that were significantly induced in control cells (log2 fold change of TE (TM/untreated) >= 1, p-adj<0.05), but with blunted induction in *eIF2A*^KO^ cells (yellow dots, Fig.4B). Translational induction of genes in response to eIF2α phosphorylation is thought to occur mainly via their 5’UTRs. Therefore, to study systematically the induction of these 12 transcripts, we cloned their 5’-UTRs into luciferase reporters. We succeeded in cloning the 5’UTRs of 11 out of the 12 transcripts. Transfection of these reporters and co-treatment with tunicamycin, however, did not show any impaired induction in *eIF2A^KO^* cells compared to control cells (Fig. 4C). This suggesting that either the induction in translation of these transcripts relies on features outside of their 5’-UTRs, or these 12 transcripts are false-positives from the ribosome footprinting.

Finally, we assessed if there is any impact of eIF2A loss on translation of uORF-containing transcripts upon tunicamycin treatment. First, we looked at the metagene profiles for footprints on all uORFs aligned to their start or stop codons, however this did not show any significant changes, apart from a minor faster transition from initiation to elongation in eIF2A^KO^ cells (Suppl. Fig. 7 A-B), which is the opposite of what one would expect. Next, we analyzed how the translation efficiency of transcripts containing either AUG- or near-cognate-initiated uORFs changes upon loss of eIF2A, compared to all transcripts, however we did not find any significant changes in either case (Suppl. Fig. 7C). Lastly, we tested the synthetic reporters containing uORFs with different start codons or Kozak sequences, described above, in the presence of tunicamycin, but found no differences in translation in eIF2A^KO^ cells compared to controls (Suppl. Fig. 7B). In sum, overall, we find a very minor, or no contribution of eIF2A on translation upon stress in HeLa cells.

## DISCUSSION

Although eIF2A was discovered prior to eIF2α, its role in translation still remains unclear, with a broad range of different functions attributed to it (reviewed in (Komar and Merrick, 2020)). In this study, we aimed to systematically assess the impact of eIF2A on translation regulation in HeLa cells. Our data show that more eIF2A is localized to the cytosol than the nucleus. Despite a previous report that eIF2A shuttles out of the nucleus in response to cellular stresses (Kim et al., 2011), we observed no change in its distribution between these compartments upon stress, consistent with previous findings in HAP1 cells (Gonzalez-Almela et al., 2018; Sanz et al., 2017). Recently, eIF2A was reported to act as a global suppressor of translation, affecting all mRNAs (Grove et al., 2023). This would result in increased global translation in *eIF2A* knock-out cells. However, we did not observe any change in bulk translation, as measured by OPP-incorporation assays, upon loss or increased expression of eIF2A. In line with no impact of eIF2A on global translation, we also did not detect any proliferation defect of eIF2A^KO^ cells, which fits with a dispensable role of eIF2A in other species (Anderson et al., 2021; Golovko et al., 2016; Kim et al., 2018; Komar et al., 2005). This is further supported by our ribosome profiling data, which showed that only a few transcripts, if any, changed in their translation upon loss of eIF2A, while the vast majority of the translatome remained unaffected. Given that eIF2A was reported to modulate translation initiation through elements in the 5’-UTR (e.g. uORFs, repeats etc.) we tested whether cloning the 5’-UTRs of affected transcripts into luciferase reporters would reveal eIF2A dependence, however, this was not the case. This suggests that if certain transcripts are affected by the lack of eIF2A this is not due to their 5’ UTRs and may rely on features in their CDS or 3’-UTRs.

The tRNA-binding properties of eIF2A remain a topic of debate. While initial studies reported that eIF2A could bind tRNA^Met^ in a GTP-independent manner (Kim et al., 2018; Zoll et al., 2002), the purification protocols used in these studies were later shown to result in eIF2D contamination, which was subsequently identified as the tRNA^Met^ binding factor, whereas eIF2A showed no such affinity (Dmitriev et al., 2010). Despite the contested tRNA-binding capabilities of eIF2A, several subsequent studies still reported a switch of tRNA delivery from eIF2α to eIF2A upon cellular stresses, when eIF2 activity is attenuated due to phosphorylation by one of the four ISR kinases (Kim et al., 2018; Kwon et al., 2017; Starck et al., 2016; Tusi et al., 2021). Taking this into consideration, we carried out ribosome profiling upon tunicamycin treatment, which results in eIF2α phosphorylation through PERK kinase. Surprisingly, also in this condition we did not find any transcript whose translation is significantly affected when comparing eIF2A^KO^ to control cells. However, we did find some transcripts whose induction was blunted in eIF2A^KO^ cells. Since no transcript showed reduced translation in eIF2A^KO^ cells compared to control cells in either the control or the tunicamycin condition, these must be transcripts which had mildly, but not significantly increased translation in the non-stressed condition and mildly, but not significantly reduced translation in the tunicamycin condition, leading to a significant difference when comparing the two treatment conditions. To further dissect if this defect in induction arises from elements within the transcript 5’-UTRs, we cloned their 5’UTRs into luciferase reporters, however, we found no significant difference between eIF2A^KO^ and control cells. Given that the more straight-forward analysis comparing translation efficiency in eIF2A^KO^ versus control cells in the tunicamycin condition revealed no transcripts with altered translation, we think the most simple explanation is that also in the stressed condition where eIF2 function is suppressed, eIF2A does not impact translation in HeLa cells.

Several reports linked eIF2A function to uORF translation. Although our ribosome profiling data indicates that there are no obvious defects in translation of transcripts with uORFs, we nonetheless decided to test the impact of eIF2A on synthetic uORF reporters. We used reporters with uORFs initiated by either AUG or by near-cognate start codons, thereby testing the role of eIF2A in leaky scanning, reinitiation, and near-cognate initiation. Our data, however, show that none of the tested reporters was affected by eIF2A loss or overexpression in both non-stressed and stressed conditions, indicating that eIF2A is dispensable for uORF translation in HeLa cell lines. We did not detect increased translation (i.e. footprints) of any other initiation factor that can potentially deliver tRNAs in *eIF2A* knock-outs (Suppl. Fig. 8), however we cannot exclude that the functional consequence of eIF2A loss is masked by another protein with redundant function.

Overall our findings are fully in agreement with recent reports showing little or no effect on translation of eIF2A loss in yeast or human HEK293-T cells (Gaikwad et al., 2024; Ichihara et al., 2021). Ichihara and colleagues generated eIF2A knockout HEK293-T cells and compared their translation to parental control cells via ribosome profiling. This revealed only 1 mRNA with reduced translation efficiency in non-stressed conditions and 4 mRNAs with reduced translation efficiency in cells treated with arsenite where eIF2 is inhibited (Ichihara et al., 2021). Thus, the lack of impact of eIF2A on mRNA translation does not appear to be specific to HeLa cells.

Historically, eIF2A was linked to translation because it was purified with other initiation factors and it shows synthetic lethality with eIF4E (Komar et al., 2005). Even though some of the initial tests with eIF2A showed no activity in reconstituted translational extracts (Merrick and Anderson, 1975), subsequent reports attributed different translational functions to eIF2A. Given that eIF2A contains an RNA-binding domain, it is possible that eIF2A plays a role in RNA trafficking or mRNA decay. The synthetic lethality between eIF2A and eIF4E (Komar et al., 2005) is interesting in this regard since eIF4E is also involved in the nuclear export of transcripts with 4E-SE elements. This raises the possibility that eIF2A might contribute to mRNA regulation beyond translation initiation. For instance, the last 50 amino acids of eIF2A are highly similar to PYM (Diem et al., 2007), a protein that binds to the 40S ribosomal subunit and to Y14, an exon-junction complex protein.

In this study we used predominantly HeLa cells, with some tests in HEK293T cells. Thus, we cannot rule out that in other cell types eIF2A might have a function in translation initiation. eIF2A knockout mice have reduced abundance of B-lymphocytes and dendritic cells in the thymic medulla, as well as lipid metabolism defects (Anderson et al., 2021). A recent study reported that mutation of *eIF2A* in Drosophila melanogaster via a MiMic transposon insertion into the second exon of *eIF2A* is lethal for the organism (Lowe and Montell, 2022). This is the first example of a lethal phenotype for eIF2A-loss in an organism, although this result needs to be confirmed with a clean knockout. The same study found that insertion of a different, piggyback transposon into the second intron resulted in viable, but infertile males due to failed sperm individualization, supporting the idea of cell-type specific function of eIF2A (Lowe and Montell, 2022). Whether these phenotypes in Drosophila are due to a role of eIF2A in translation initiation, however, remains to be investigated. Overall our results support the idea that eIF2A plays a minor, or no role in regulating translation initiation in human HeLa and HEK293T cells.

## MATERIALS & METHODS

### Cell lines, culture conditions, and treatments

Cell lines were cultured in DMEM (Gibco 41965039) supplemented with 10% fetal bovine serum (FBS) (Sigma, S0615) and 100 U/ml Penicillin/Streptomycin (Gibco 15140122). Cell splitting was done with a quick PBS wash and treatment with Trypsin-EDTA (Gibco, 25200056). All cell lines were tested negative for mycoplasma and authenticated using SNP typing. For the indicated experiments, cells were treated either with DMSO, or with 100 µM Sodium arsenite (Sigma, S7400-100G), 1µg/ml poly (I:C) (Tocris Bioscience, 4287/10), 1 µg/ml lipopolysaccharides LPC (Sigma, L2630) or 1 µg/ml tunicamycin (PanReac AppliChem, A2242,0005) for the indicated periods of time. For siRNA mediated knock-down, cells were reverse-transfected during seeding with 1.5 µl of 20 µM siRNA mix and 9 µl Lipofectamine RNAiMax reagent (Invitrogene 13778075). 72 hours post transfection, cells were re-seeded in 96-well format to perform luciferase assays. Sequences of siRNAs used in this study are provided in Suppl. Table 1.

### Generation of knockout cell lines and targeted CRISPR-Cas9 screen

eIF2A knock-out HeLa and HEK293T-H-Kb cell lines were generated using CRISPR-Cas9 with sgRNA sequences designed using CHOPCHOP software (Labun et al., 2019), and listed in Supp. Table 2. Oligos coding for the sgRNA sequences were cloned into pX459V2.0 (Doench et al., 2016) via the Bbs1 site, then wildtype HeLa or HEK293T-H-Kb cells were transfected with the plasmids using Lipofectamine 2000 in a ratio of 2:1 reagent:DNA (Life Technologies, 11668500). 24 hours post-transfection, transfected cells were selected with medium containing 1.5 µg/ml puromycin (Sigma-Aldrich, P9620). After three days, surviving cells were shifted into normal medium (1x DMEM, 10% fetal bovine serum, 1% Penicillin/Streptomycin) and regrown to confluence. Single clones were selected by serial dilution into 96 well plates. Loss of protein in expanded single clones was tested with anti eIF2A antibodies by immunoblotting and clones were confirmed by genotyping.

### Preparation of cell lysates with RIPA buffer

Cells were seeded at a density of 500.000 cells per 6-well. Following a treatment, cells were washed briefly with PBS, and then lysed with 120µl of RIPA buffer supplemented with 20U of Benzonase (Merk Millipore, 70746-3), protease (Sigma, 4693159001) and phosphatase (Sigma, 4906837001) inhibitors. Cells were collected by scraping and the lysate was clarified by centrifugation at 4°C for 10 minutes at 20.000g. The protein concentration was measured by Pierce BCA (Life technologies, 23224, 23228). Samples were balanced to equal protein concentration and mixed with 5x Laemmli buffer (1/5 of the final volume). Samples were incubated at 95°C for 5 minutes and then loaded on an SDS-PAGE gel.

### Western blotting

Cell lysates were separated on SDS-PAGE gels, and transferred to a nitrocellulose membrane with 0.4 µm pore size (Amersham, 10600002) by wet transfer. To block unspecific binding, membranes were blocked in 5% skim milk / PBST for 1 hour. Membranes were then probed by overnight incubation in primary antibody solution (5% BSA / PBST) at 4°C. On the following day membranes were washed three times 15’ in PBST and incubated in secondary antibody (1:10000 in 5% skim milk / PBST) for 2 hours at room temperature. To remove unbound secondary antibodies, membranes were washed three times for 15 minutes in PBST. Finally, chemiluminescence was detected with ECL reagents (Thermo Schientific, 32109) and imaged with a Biorad ChemiDoc imaging system. Antibodies used for immunoblotting are listed in Suppl. Table 3.

### Cloning

Firefly (pAT1620) and Renilla luciferase (pAT1618) under control of LMNB1 5’UTR were described previously (Schleich et al., 2017). The 5’UTRs of eIF2A-dependent candidate transcripts, as well as integrated stress responsive 5’UTRs, were PCR amplified from HeLa cDNA with the oligos indicated in Suppl. Table 4. PCR products were gel purified, digested with HindIII and Bsp119I, and subsequently cloned into pAT1618 via the same sites. If the 5’ UTR length was below 120 nucleotides, it was directly oligo cloned into pAT1618 via HindIII and Bsp119i. The cloned 5’ UTR variants are listed in Suppl. Table 5 with the indicated transcript ID. Reporters with uORFs with different Kozak strengths were generated and described previously (Bohlen et al., 2023). The reporters with uORFs starting with near cognate start codons were generated for this study by substituting the AUG-uORF generated in (Bohlen et al., 2023) with the near-cognate one via oligo-cloning using the Kpn2I and BshT1 sites with oligos listed in Suppl. Table 4. The EMCV-IRES reporter was generated and described previously in (Roiuk et al., 2024). The short 5’-UTR reporter was produced by oligocloning via SacI and Bsp119I sites in pAT1618. All generated constructs were verified by sanger sequencing.

### Ribosome profiling

Control or eIF2^KO^ HeLa cells were seeded at 1.2 million cells per 15 cm dish in 20 ml of growth medium two days before harvesting. The following day, cells were treated either with DMSO or 1 µg/ml tunicamycin for 16 hours. After treatment, cells were quickly washed with ice-cold 1xPBS supplemented with 10 mM MgCl2 and 800 µM Cycloheximide. After the wash, all residual solution was removed by gentle taping of the 15cm dish on its side, followed by cell lysis with 150 µl of the following buffer: 0,25 M HEPES pH 7.5, 50 mM MgCl2, 1 M KCl, 5% NP40, 1000 μM Cycloheximide. Cells were scraped into an Eppendorf tube and the lysate was clarified by centrifugation at 15.000xg for 10 minutes at 4°C. The concentration of lysate was estimated using a nanodrop spectrophotometer, measuring RNA content against a water-blanked control. 150 ul of lysate was used for the total RNA preparation with the RNeasy kit (Qiagen, cat. No. 74106). The remaining lysate was used for treatment with RNaseI (100 Unit per 120µg of lysate) on ice for 30 minutes. Following digestion, the lysates were loaded on a 15-65% sucrose gradient, which was prepared in advance with the use of a Biocomp Gradient Master. The lysate was ultracentrifuged for 3 hours at 35000 rpm in a Beckman Ultracentrifuge with a SW40Ti rotor. To collect the 80S fraction the gradient was separated on a Biocomp Gradient Profiler system. The collected 80S fractions were used for RNA extraction with acid-phenol. Briefly, the volume of the sample was adjusted to 700µl with 10mM Tris pH 7.0 and mixed with 750 µl of prewarmed acid phenol. Sample-phenol mix was incubated at 65C with constant shaking at 1400 rpm for 15 minutes, followed by incubation on ice for 5 minutes. The sample was subsequently centrifuged at 20000g for 2 minutes and the supernatant was transferred into a new tube, containing 700µl of acid phenol. After 5 minutes of incubation at room temperature, the sample was spun at 20000g for 2 minutes and the supernatant was transferred into the tube with 600 µl of chloroform. The sample was then mixed by vortexing and centrifuged at 20000g for 2 minutes. The supernatant fraction was transferred into a new tube and mixed with equal volume of isopropanol, 2µl of Glycoblue (Invitrogene AM9516) and 1/10 volume 3M NaAc pH5.2. To precipitate RNA, the sample was incubated overnight at −80C. The following day, the sample was centrifuged at 4C for 30 minutes at 14000 rpm, followed by a 70% Ethanol wash. The RNA-pellet was resuspended in RNase-DNase free water. The integrity of all RNA samples was analysed on a Bioanalyser. To size-select footprints, RNA extracted from the 80S peak was run on a 15% Urea-Polyacrylamide gel and fragments of 25-35 nucleotides were purified from the gel. For this, the gel pieces containing the footprints were broken into small pieces with gel smasher tubes. 0.5 ml of 10 mM Tris pH 7 were added to the smashed gel pieces and the suspension was incubated at 70°C for 10 minutes with shaking. The mix was briefly centrifuged and the supernatant was used for RNA precipitation by isopropanol. Purified footprints were phosphorylated by means of T4 PNK (NEB) for 1 hours at 37°C in PNK buffer supplemented with 10 mM ATP. After this, the footprints were again precipitated and purified using isopropanol. To estimate the quality of footprints, RNA was run on an Agilent Bioanalyzer small RNA chip, followed by library preparation with the Next-Flex small RNA v.4 kit protocol (Perkin Elmer, NOVA-5132-06), in accordance to manufacture recommendations. Total RNA libraries were prepared using the Illumina TruSeq Stranded library preparation kit. The quality of libraries were checked on a Bioanalyser with the use of a High sensitivity DNA kit (Agilent, 5067-4626). The libraries were sequenced on an Illumina Next-Seq 550 system.

### Data Analyses of Ribosome profiling

Reads were trimmed from adaptors and randomized nucleotides derived from use of the Nextflex kit with cutadapt software. By use of bowtie2 the reads aligning to tRNA or rRNA were removed. All remaining reads were mapped to the human transcriptome (Ensemble transcript assembly 94) and genome (hg38) using BBMap software, with multiple mapping allowed. The reads mapping to the coding sequences were quantified with lab-based software written in C (https://github.com/aurelioteleman/Teleman-Lab). For each transcript the value of reads per kilobase of coding sequence was estimated and only transcripts with values more or equal of 2.5 were used for the subsequent analysis. Metagene profiles were built with the custom-made software written in C. For the metagene profile of uORF stop codons, only transcripts with a space of more than 50 nucleotides between uORF and main ORF were used. The DESeq2 software package was used to calculate log2 fold-changes and adjusted p values for the difference in translation efficiency (defined as ribosome footprints / total RNA) in control versus eIF2A knockout cells. DESeq2 was run with the design = ∼assay + condition + assay:condition.

### OPP incorporation assay

A total of 0.5 million control or eIF2A^KO^ HeLa cells were seeded in six-well plates a day prior to treatment. On the next day, cells were labelled by incubation with 20 μM OPP reagent (Jena Bioscience NU-931-05) for 30 min. Negative control sample was pre-treated with 100µg/ml cycloheximide, and the cycloheximide was maintained during OPP labelling. Following labelling, the cells were lysed, the concentration of samples was measured and equalized, and the incorporation of OPP was estimated via western blot using anti-puromycin antibodies.

### Dual-luciferase translation reporter assay

Cells were seeded in 96-well plate format at a density of 8000 cells per well. The following day, cells were transfected with Lipofectamine 2000 with 100 ng of Renilla luciferase plasmid and 100 ng of Firefly luciferase plasmid per well. In case of tunicamycin (TM) treatment, 6 hours post transfection, medium was exchanged with fresh medium supplemented either with either DMSO or 1µg/ml TM. After 16 hours the luciferase assay was carried out using the Promega Dual-Luciferase assay system (Promega, E1910) following manufacturer’s instructions.

### Immunofluorescent detection of overexpressed Flag-eIF2A

15,000 control HeLa cells were seeded in 12-well format on poly-lysine coated 12-mm cover slips. The following day, cells were transfected with plasmid coding for eIF2A-Flag using Lipofectamine 2000 in a ratio 1:2 DNA:reagent (Life Technologies, 11668500). One day later, cells were treated with or without sodium arsenite (100µM, 1h) and then washed with PBS and fixed by incubation for 15 minutes in 4% formaldehyde in PBS. Cells were permeabilized by incubation for 10 minutes in 0.2% Triton-X-100 in PBS and blocked for 1 hour in 0.25% BSA in PBS. Following blocking, cells were incubated overnight at 4C in primary antibodies. On the following day, the cells were washed twice with blocking buffer and incubated with fluorophore-conjugated secondary antibodies for 1 hour. Finally, cells were shortly stained with 5 µg DAPI (Applichem A1001) and mounted on glass slides using Vectashield (Vector Labs H-1000). The distribution of Flag-eIF2A was analysed on a confocal microscope (Leika TCS SP8) using a 63x objective.

### Cell proliferation assay

For cell proliferation assays, cells were seeded into 96 well plates at a density of 1000 cells per well in 100 µl of medium, 8 wells per condition. A total of 5 plates were seeded for the sequential sample collection. The proliferation curve was built by harvesting one plate every 24 hours and performing the Cell Titer Glo (Promega, G7572) according to the manufacturer’s instructions.

### Subcellular fractionation

On the day prior to the experiment, cells were seeded in 10cm dish format, 1.5-2 million cells per dish. The next day, cells were treated for 2 hours either with 1µg/ml Tunicamycin or DMSO, followed by a quick wash with PBS and harvesting by trypsinization. The collected cells were pelleted by gentle centrifugation at 4C (3000g 5 minutes). The supernatant was removed, and the cell pellet was quickly washed twice with PBS. Cells were resuspended with 200µl of 1x Hypotonic Buffer (20mM Tric-HCl, pH7,4; 10mM NaCl, 3mM MgCl) by gently pipetting up and down. The resuspended cells were incubated on ice for 15 minutes, followed by addition of 1/20 of the suspension volume of 10% NP40. Cells with NP40 were vortexed for 10 seconds at highest speed and centrifuged at 3000g for 10 minutes at 4C. The supernatant was moved into a new tube and marked as the cytosolic fraction, while the pellet was washed twice with PBS, moved into a new tube and resuspended in 200 ul of Cell Extraction Buffer (10mM Tris,pH 7.4, 2mM Na_3_VO_4_, 100mM NaCL, 1% Triton X-100, 1mM EDTA, 10% glycerol, 1mM EGTA, 0.1% SDS. 1mM NaF, 0.5% deoxycholate, 20mM Na_4_P_2_O_7_). The pellet was incubated on ice for 30 minutes with occasional vortexing, followed by centrifugation at 14000g at 4C for 30 minutes. The supernatant was collected and marked as the nuclear fraction.

*Simultaneous detection of a uORF/oORF peptide and mainORF mNeonGreen* The HEK293T-K^B^ *eIF2A*^KO^ cell line was generated the same way as HeLa-*eIF2A*^KO^ cells. Two days prior to the experiment, cells were seeded in 6 well format. The following day, cells were transfected with empty vector, or with plasmids encoding mNeonGreen-PEST carrying either an oORF-less 5’UTR, or a 5’UTR containing a uORF coding for “SIINFEKL” with either AUG or near-cognate start codons. The AUG codon was placed in a Kozak context predicted to be either ‘medium’ or ‘strong’. 24 hours post-transfection, cells were washed with PBS, collected by trypsinization and resuspended and incubated for 10 minutes in blocking buffer (1% BSA in 1xPBS). After blocking, cells were washed once more with PBS and stained for 30 minutes in the dark at 4C with 0.25 ug/ml monoclonal Antibody OVA257-264, which detects the SIINFEKL-peptide bound to H-2Kb (Life technologies 25-5743-82). Following staining, cells were washed three times with blocking buffer to remove unbound antibodies, resuspended with 300 ul of blocking buffer and analysed with a Guava® easyCyteTM flow cytometer running Guava Soft 3.3

### Quantitative RT-PCR

Total RNA from either control of eIF2A^KO^ HeLa cells was extracted using RNase Mini spin columns (Qiagen, cat. no. 74106). Reverse transcription (RT) of 1ug of total RNA with random hexamer and oligo-dT+ primers using Maxima H minus reverse transcriptase was performed to generate cDNA. The amplification efficiency of all Q-RT-PCR primer pairs was checked using serial dilution of a sample. Quantitative RT-PCR was run on a QuantStudio3 instrument with primaQUANT SYBRGreen low ROX master mix. RNA levels were normalized to the levels of either *GAPDH* or *RPL13A* mRNA. Sequences of oligos used for Q-RT-PCR are provided in Supplemental Table 6.

## ACKNOWLEDGEMENTS

We thank the DKFZ Genomics Core Facility for next-generation DNA sequencing.

---

**Suppl. Figure 1:**
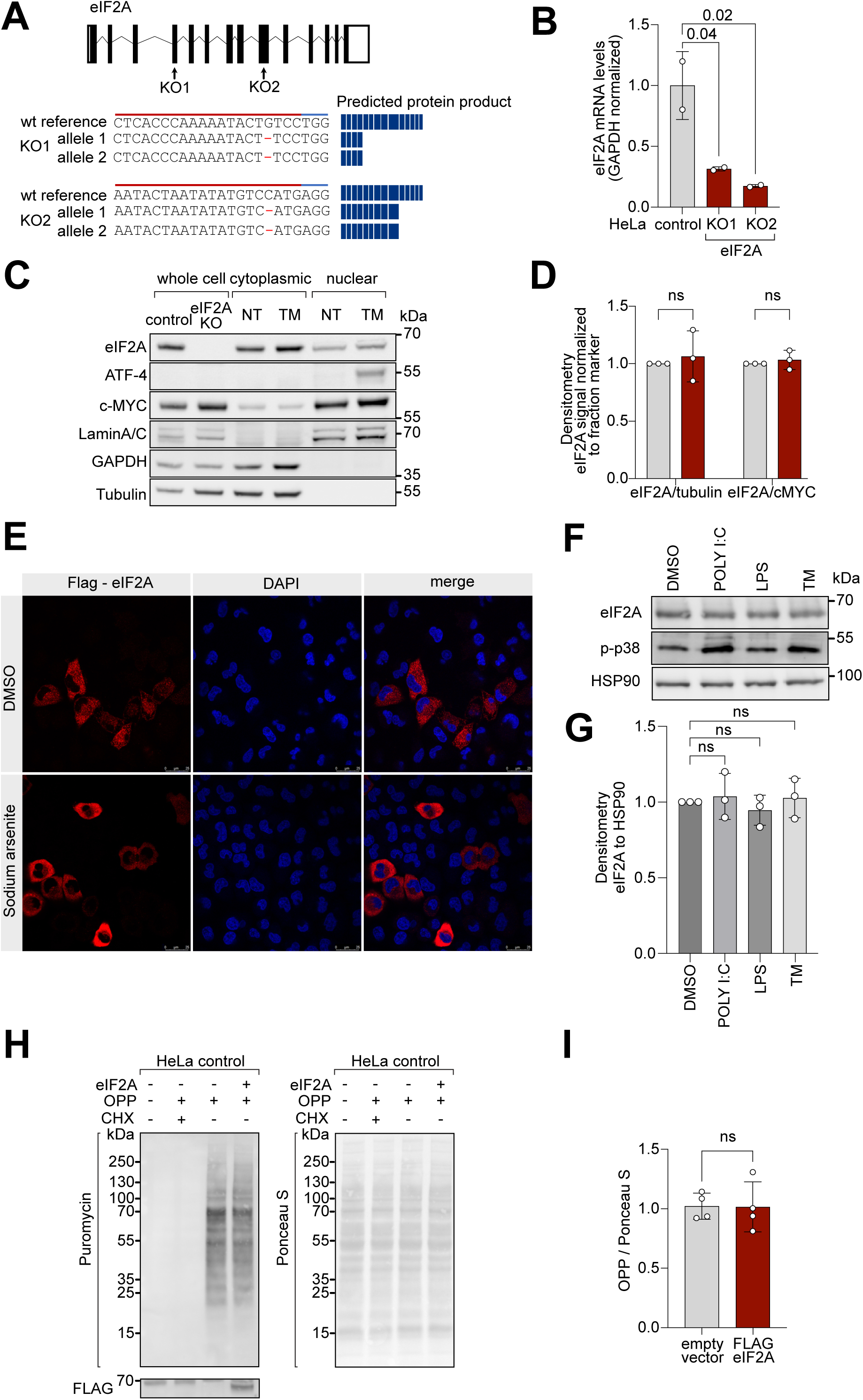
**Loss of eIF2A does not perturb cell proliferation and global translation.** (**A**) Genotyping of *eIF2A*^KO^ cell lines. Two eIF2A knockout lines were generated by targeting two separate coding exons. Shown is the resulting mutation at the DNA level, and a schematic of the predicted protein product. (**B**) mRNA levels of *eIF2A,* quantified by qRT-PCR, are reduced in the eIF2A^KO^ lines. Error bars: standard deviation. Significance by ANOVA with Dunnett’s multiple comparison test. **(C-D)** Subcellular localization of eIF2A, detected by immunoblotting cytosolic and nuclear fractions or whole cell lysate (WCL), does not change upon treatment with tunicamycin (TM). Cells were treated with DMSO or Tunicamycin (1 µg/µl) for 2 hours. (C). Representative immunoblot (C) of three independent replicates quantified in (D). Nuclear and cytosolic fractions were lysed in equal volumes to reflect the relative contribution of the two compartments in a cell. Error bars: deviation. Significance by multiple unpaired, t-test. (**E**) Overexpressed FLAG-eIF2A shows predominantly cytoplasmic localization, assessed by immunofluorescent staining with anti-FLAG antibodies. Cells transfected with plasmid expressing eIF2A-FLAG were treated for 1 hour either with DMSO or 100 µM Sodium arsenite (SA). **(F-G)** eIF2A levels in HeLa cells are not affected by poly (I:C) (1µg/ml), LPC (1 µg/ml) or tunicamycin (1µg/ml) for 3 hours. Phospho-p38 signal is used as a positive control for the treatment. Representative immunoblot (F) or three independent replicates quantified in (G). Error bars: standard deviation. Significance by ANOVA with Dunnett’s multiple comparison test. **(H-I)** Global protein translation levels in HeLa cells do not change upon Flag-eIF2A overexpression, assessed by immunoblot against OPP (left) and normalized to total protein amount assayed by PonceauS (right). Representative blots in (H), of four independent replicates quantified in (I). Error bars: standard deviation. Significance by unpaired, two-sided, t-test. All panels: ns = not significant.

**Suppl. Figure 2:**
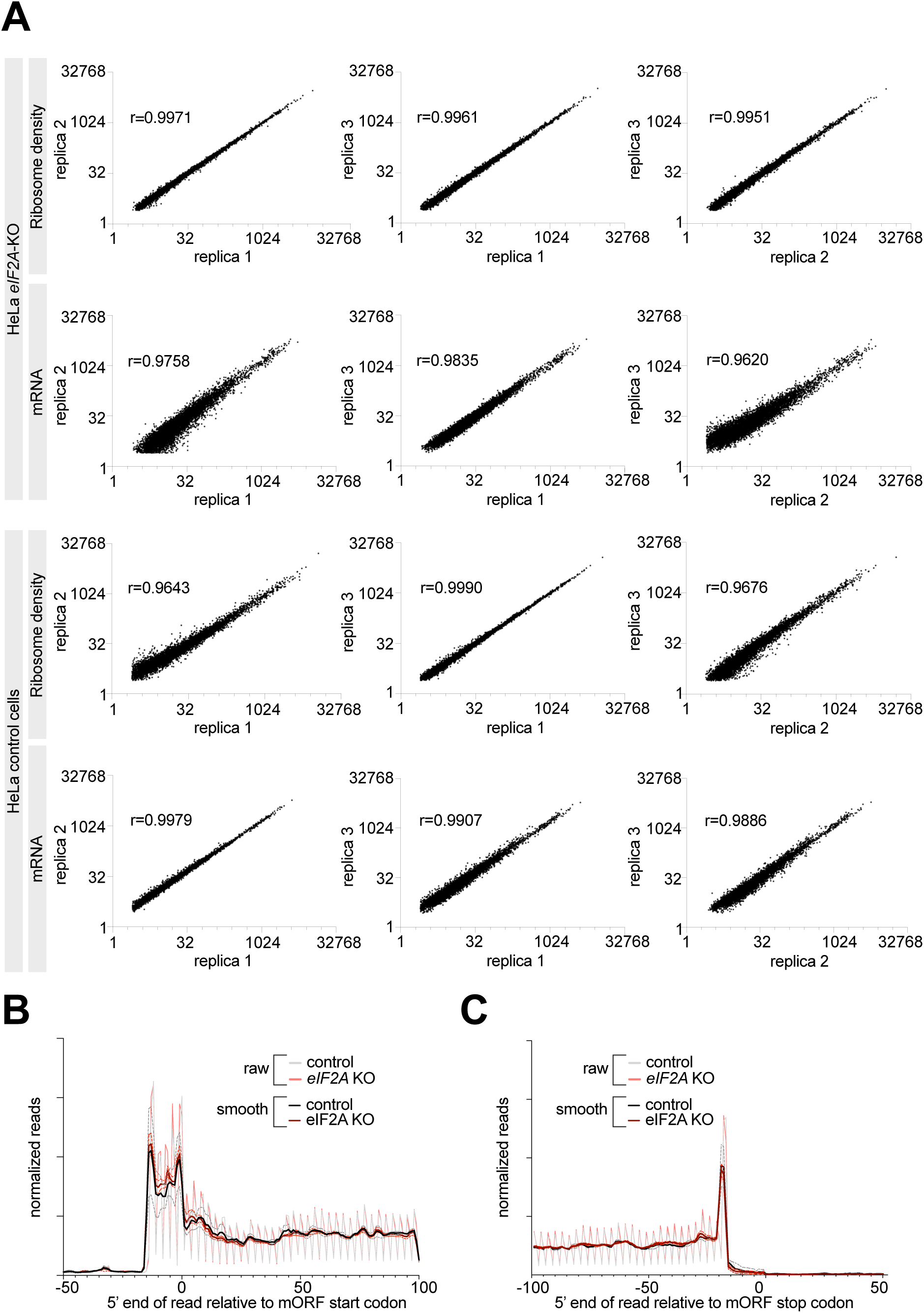
Ribosome profiling of control and eIF2A^KO^ HeLa cells. **(A)** Reproducibility between replicates of Ribosome profiling and total-mRNA libraries is shown. Three biological replicates were generated for control and eIF2A^KO^ HeLa cells each. The Pearson’s coefficient (r) is shown for each compared pair. **(B-C)** Metagene profiles of footprints aligned to either the start codon (B) or the stop codon (C) of all main Open Reading Frames, for control and eIF2A^KO^ HeLa cells. “Smooth” curves were generated by averaging read counts with the sliding window of 3 nt. The dotted lines indicated standard deviation between three replicates.

**Suppl. Figure 3:**
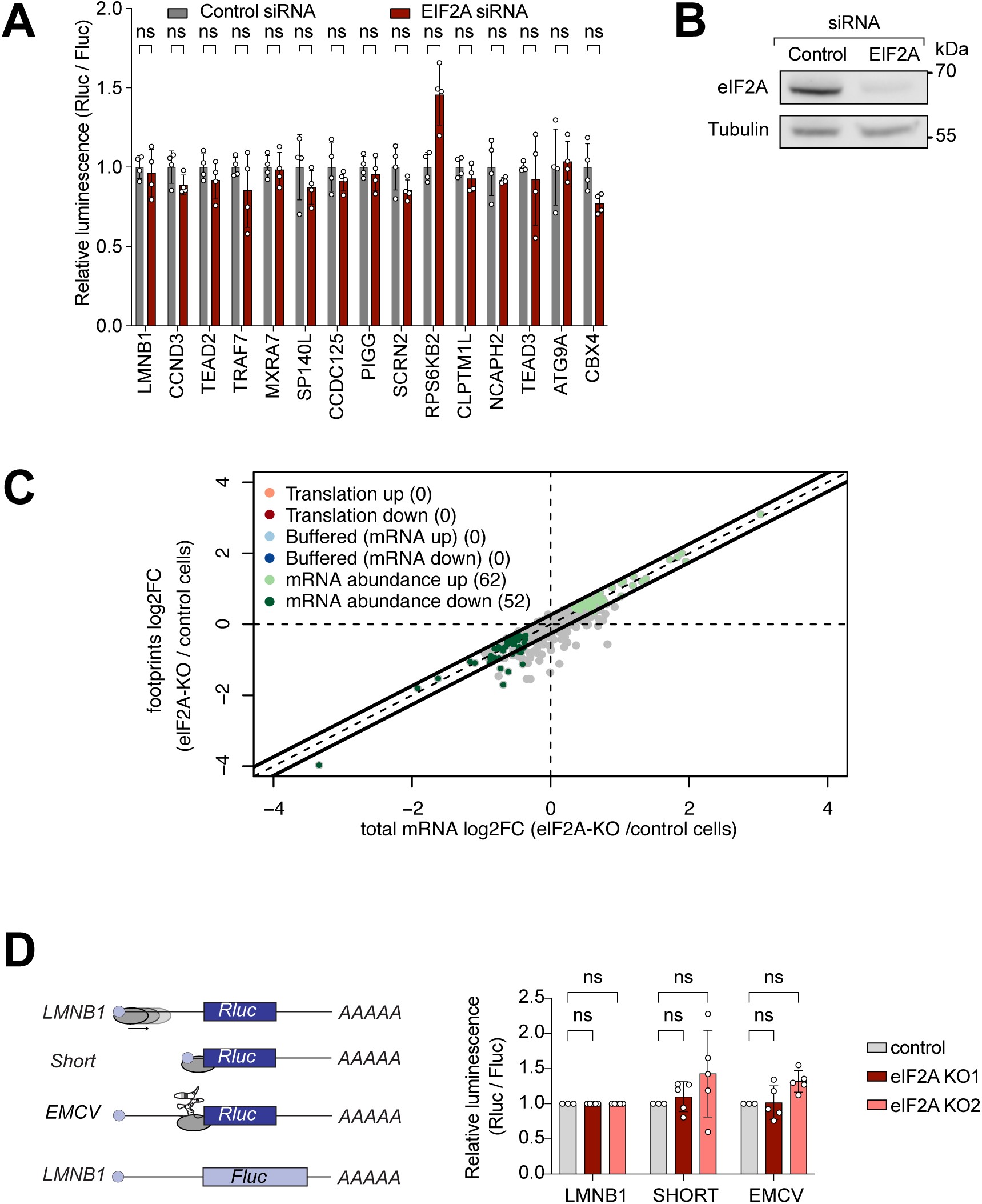
Loss of eIF2A does not affect translation of multiple different types of reporters. **(A)** Luciferase reporters harboring 5’ UTRs of transcripts predicted to be eIF2A-dependent from ribosome footprinting, do not show significantly reduced translation upon siRNA mediated knockdown of *eIF2A*. The 5’ UTRs of the indicated genes were cloned upstream of Renilla Luciferase (RLuc) and co-transfected with a Firefly Luciferase (FLuc) normalization control reporter. Negative control RLuc reporter and the FLuc normalization control carry the 5’UTR of lamin B1 (LMNB1). Significance by multiple unpaired t-test, ns = not significant. Error bars represent standard deviation. **(B)** Western blot control for efficiency of siRNA-mediated knockdown of *eIF2A*. **(C)** Transcripts shown in Fig. 2A were reanalyzed with anota2seq package (Oertlin et al., 2019). Scatter plot of log2 fold-change of total RNA eIF2A^KO^/ control (x axis) versus log2 fold-change of footprints eIF2A^KO^/ control (y axis) is shown. Significant changes are detected in mRNA levels, while loss of eIF2A does not perturb translation. **(D)** Transfection of luciferase reporters designed to place the ribosome directly on top of the initiation AUG do not show reduced translation in eIF2A^KO^ cells compared to controls. A 5’UTR containing the EMCV IRES or a short 5’ UTR of only 12 nt was cloned upstream of Renilla Luciferase (RLuc) and co-transfected with a Firefly Luciferase (FLuc) normalization control. Negative control RLuc reporter and the FLuc normalization control carry the 5’UTR of Lamin B1 (LMNB1). Significance by ANOVA with Dunnett’s multiple comparison test. error bar = st. dev., ns = not significant.

**Suppl. Figure 4:**
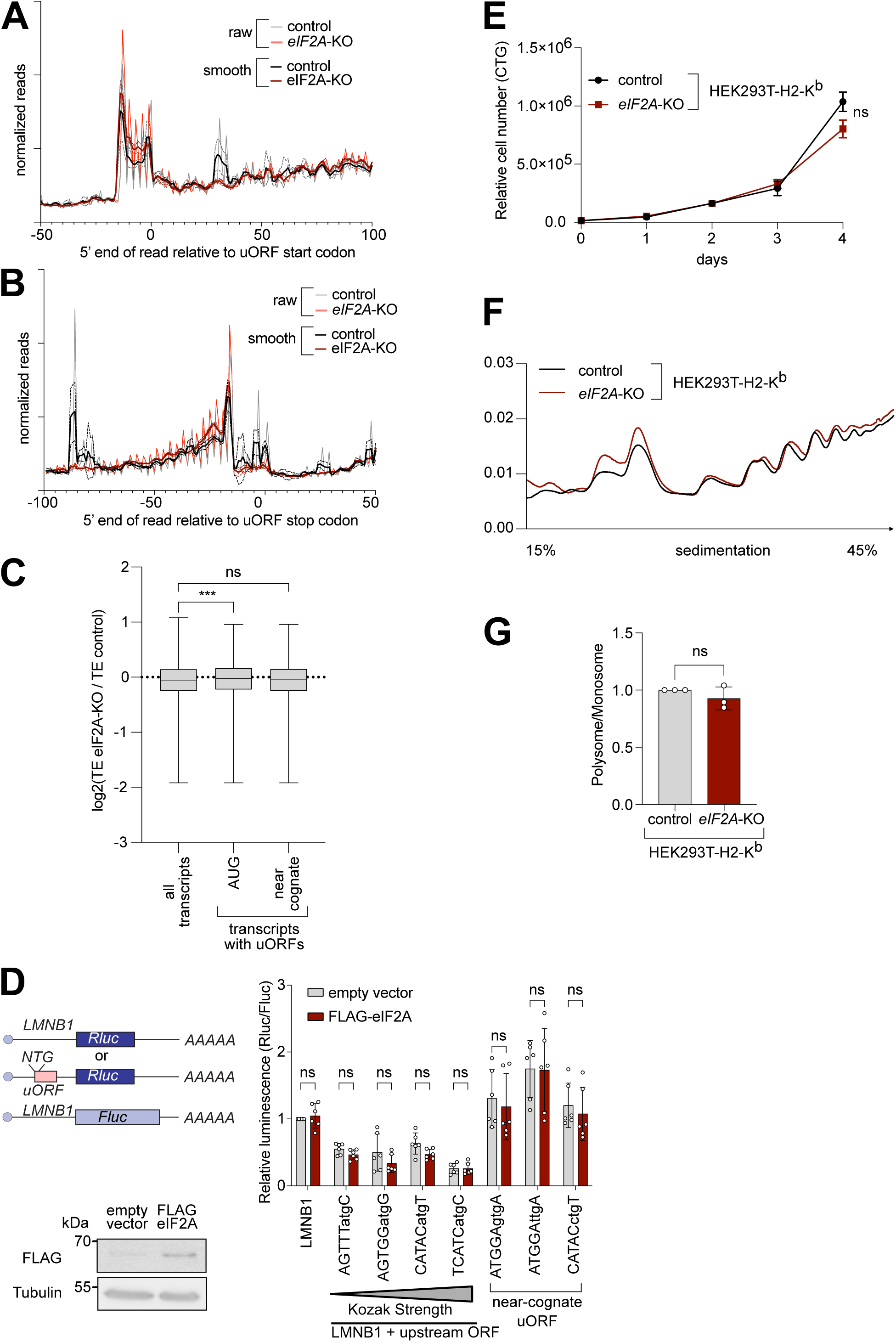
Knockout of *eIF2A* has no effect on uORF translation. **(A-B)** Metagene profiles of footprints relative to either the start codon (A) or stop codon (B) of uORFs transcriptome-wide, for control or eIF2A^KO^ HeLa cells. “Smooth” curves were generated by averaging footprint counts with a sliding window of 3nt. The dotted lines indicated standard deviation between three replicates. (**C**) eIF2A has little impact genome wide on translation of mRNAs with uORFs. Comparison of the change in translation efficiency between eIF2A^KO^and control HeLa cells for three different groups of mRNAs: all transcripts, transcripts with AUG-initiated uORFs, or transcripts with uORFs that start with a near-cognate codon. Significance by ANOVA with Dunnett’s multiple comparison test. ns = not significant, *** p<0.001 (**D**) Activity of synthetic reporters harboring uORFs with different start codons and initiation contexts is not altered upon eIF2A-overexpression. The sequence context of the uORF start codons is indicated: either AUG or a near-cognate start codon (GTG, TTG, CTG) was used. Overexpression of eIF2A was validated by western blot against the FLAG-tag. Significance by Dunnett’s multiple comparison test ANOVA, error bar = st. dev. ns = not significant. (**E**) Proliferation of eIF2A^KO^ or control HEK293T-H2-K^b^ cells by CellTiter Glo. Error bars: standard deviation. Significance by unpaired, two-sided, t-test. ns = not significant. **(F-G)** Global cellular translation, assayed via polysome profiles, shows little difference between eIF2A^KO^ and control HEK293T-H2-K^b^ cells. Lysates from either eIF2A^KO^ or control HEK293T-H2-K^b^ cells were separated on sucrose gradients. One representative graph is shown in (F). The polysome/80S ratio of three independent replicates is shown in (G). Error bars: standard deviation. Significance by unpaired, two-sided, t-test. ns = not significant.

**Suppl. Figure 5:**
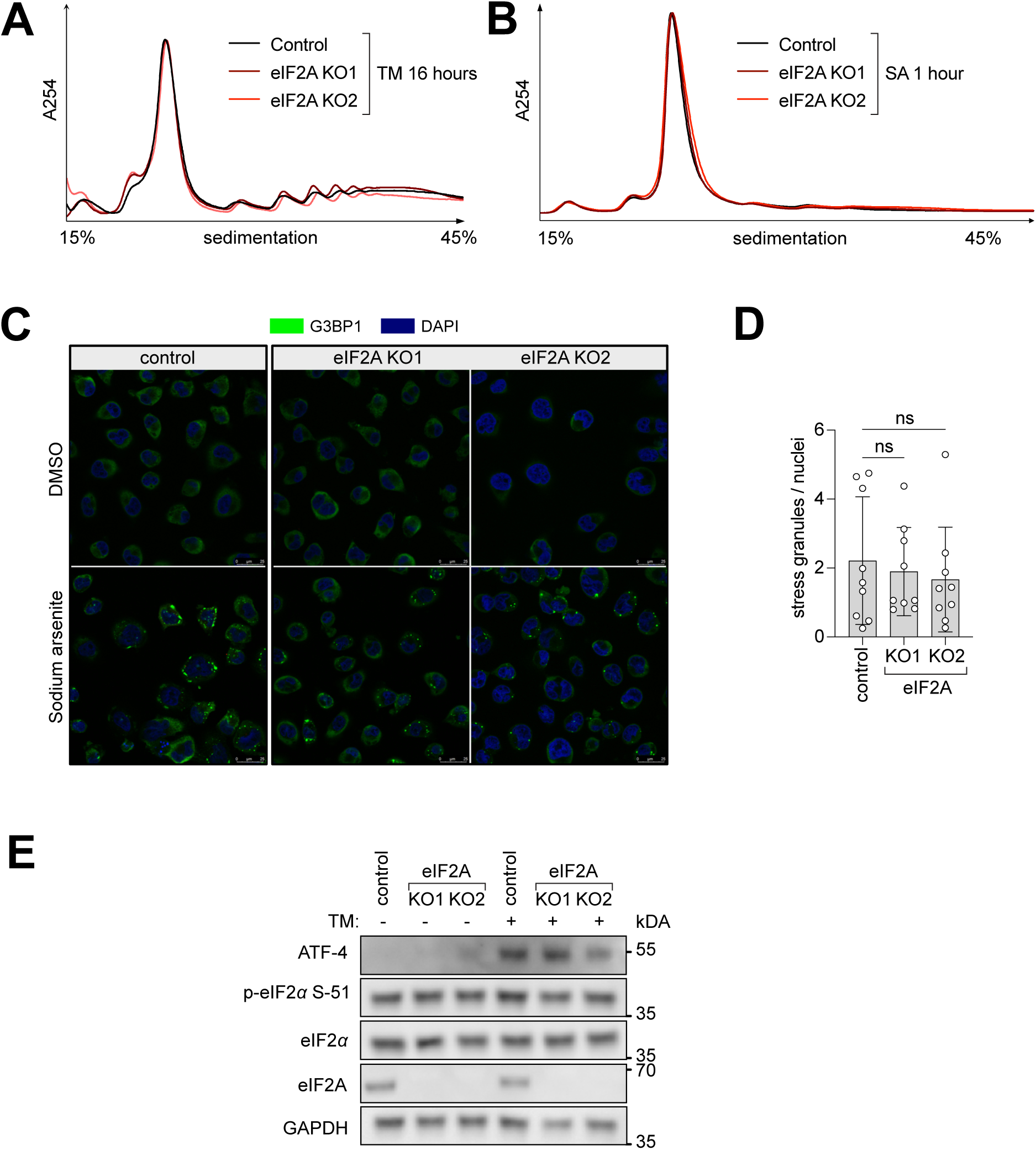
The Integrated Stress Response suppresses translation equally well in control and *eIF2A*^KO^ HeLa cells. **(A-B)** Polysome profiles of eIF2A^KO^ versus control HeLa cell lines show hardly any differences upon stress caused either with (A) 1 µg/ml tunicamycin (TM) for 16 hours or (B) 100µM sodium arsenite (SA) for 1 hour. Profiles were aligned by the height of the 80S peak. **(C-D)** eIF2A knockout cells form stress granules to the same degree as the parental control HeLa cell line. Control or eIF2A^KO^ HeLa cells were treated for 2 hours with 100 µM sodium arsenite and stained for G3BP1. (C) Representative images. (D) The number of stress granules, normalised to the number of nuclei in each field of view is shown. Significance by ANOVA with Dunnett’s multiple comparison test. ns = not significant. **(E)** *eIF2A*^KO^ and control HeLa cells phosphorylate eIF2α to the same degree in response to tunicamycin (1µg/mL for 16 hours). ATF-4 is used as a marker for downstream activation of the integrated stress response.

**Suppl. Figure 6:**
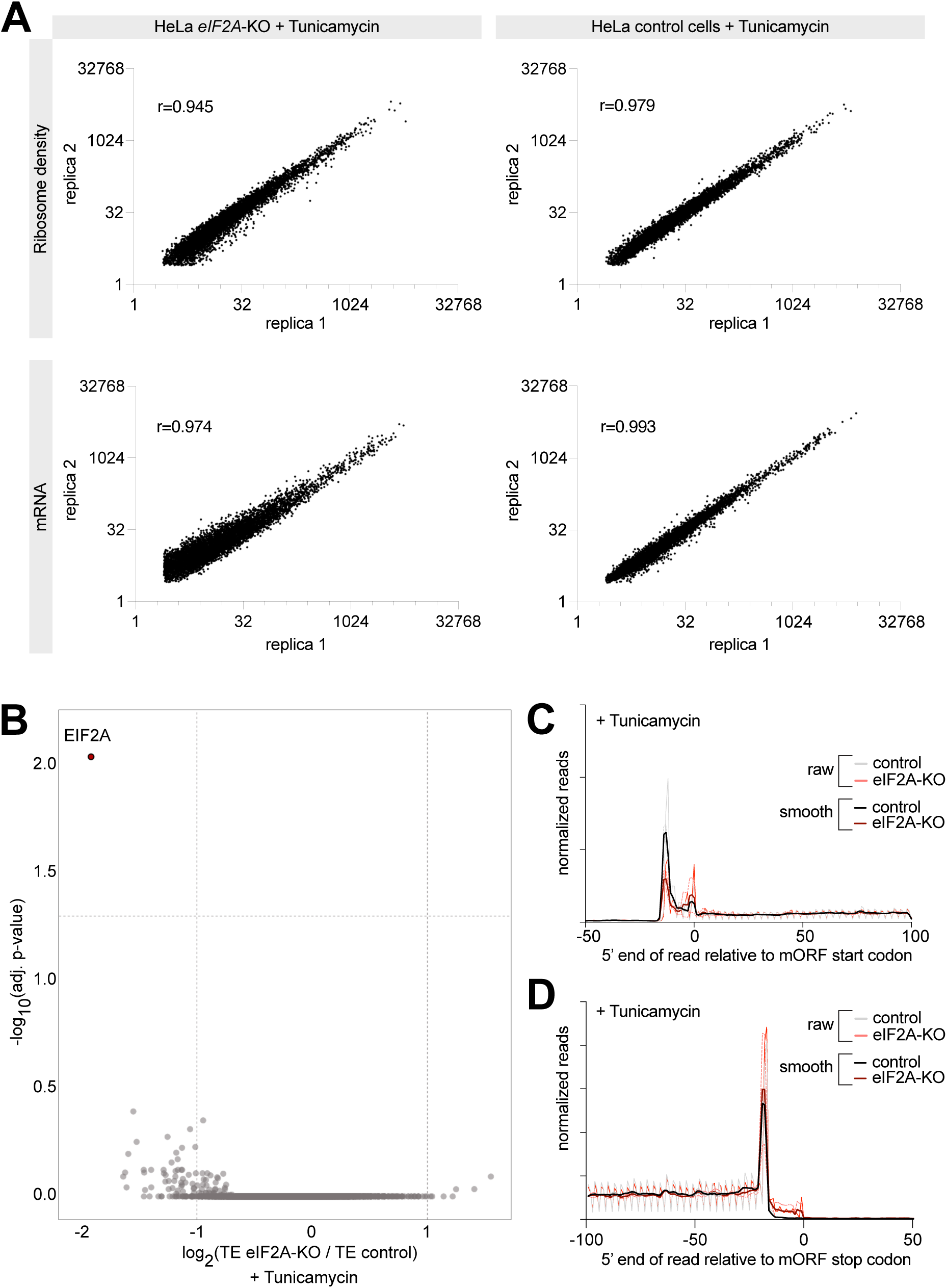
Ribosome profiling of tunicamycin treated *eIF2A*^KO^ and control HeLa cells. (**A**) Reproducibility between replicates of Ribosome profiling or total-mRNA libraries from eIF2A^KO^ or control HeLa cells treated with tunicamycin. Two biological replicates were generated for each genotype. The Pearson’s coefficient (r) is shown for each comparison. (**B**) Ribosome profiling identifies zero transcripts affected by eIF2A knockout in tunicamycin treated condition. Scatter plot of log2(fold change) of Translation Efficiency comparing eIF2A^KO^ to control HeLa cells, both treated with tunicamycin, on the x-axis, versus significance on the y-axis. Significant candidates with log2(fold change) < −1 are shown in red. Significance was estimated with the Wald test performed by DESeq2 package. p-values are adjusted for multiple comparison. **(C-D)** Metagene profiles of footprints aligned to either the start codon (C) or the stop codon (D) of all main Open Reading Frames, for control and eIF2A^KO^ HeLa cells, both treated with tunicamycin. “Smooth” curves were generated by averaging read counts with the sliding window of 3 nt. The dotted lines indicate standard deviation between three replicates.

**Suppl. Figure 7:**
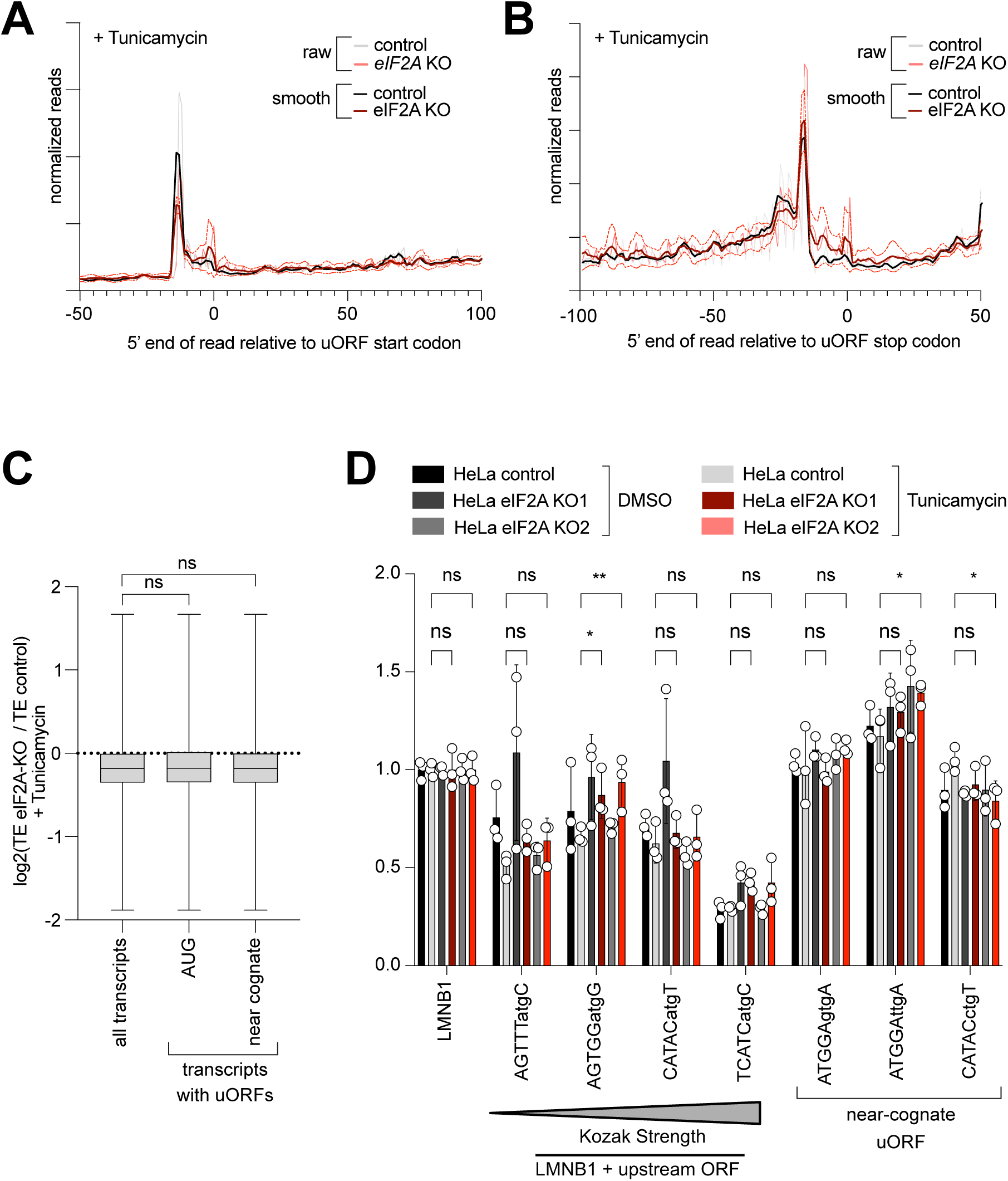
Translation of uORF-bearing transcripts is not affected upon loss of eIF2A in tunicamycin treated cells. **(A-B)** Metagene profiles of footprints relative to either the start codon (A) or stop codon (B) of uORFs transcriptome-wide, for control or eIF2A^KO^ HeLa cells, both treated with tunicamycin. “Smooth” curves were generated by averaging footprint counts with a sliding window of 3nt. The dotted lines indicated standard deviation between three replicates. (**C**) eIF2A has little impact genome wide on translation of mRNAs with uORFs in cells treated with tunicamycin. Comparison of the change in translation efficiency between eIF2A^KO^ and control HeLa cells, both treated with tunicamycin, for three different groups of mRNAs: all transcripts, transcripts with AUG-initiated uORFs, or transcripts with uORFs that start with a near-cognate codon. Significance by ANOVA with Dunnett’s multiple comparison test. ns = not significant. (**D**) Synthetic reporters harboring uORFs with different start codons and initiation contexts do not show dependence on eIF2A also when the integrated stress response is activated, thereby suppressing eIF2 function. Either control or eIF2A^KO^ HeLa cells were transfected with the indicated reporters and then treated with 1µg/mL tunicamycin for 16 hours prior to assaying luciferase activity. The sequence context of the uORF start codons is indicated: either AUG or a near-cognate start codon (GTG, TTG, CTG) was used. Significance by Dunnett’s multiple comparison test ANOVA, error bar = st. dev. ns = not significant. * p < 0.05, ** p < 0.01.

**Suppl. Figure 8:**
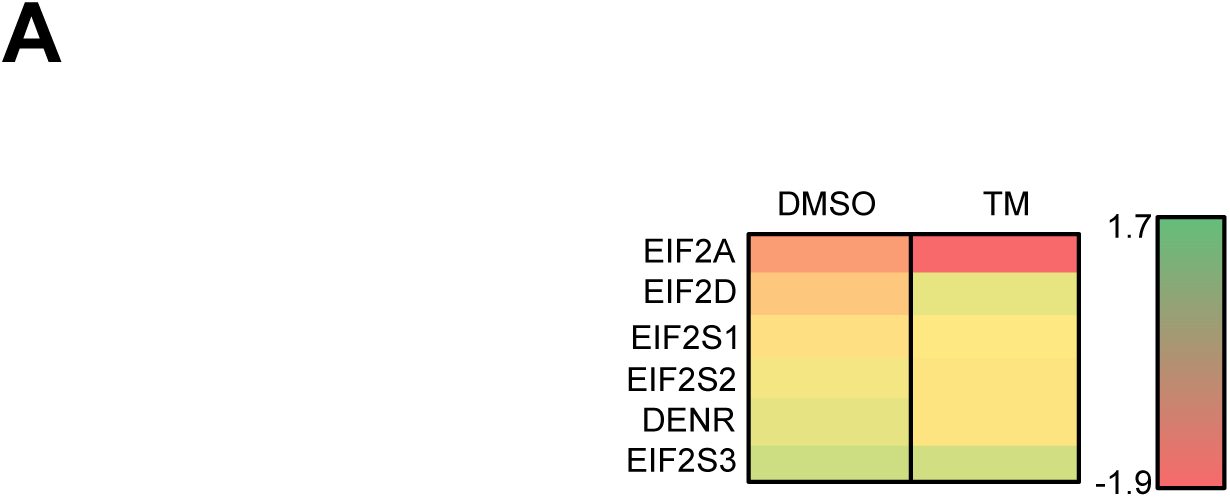
Expression of initiation factors reported to possess tRNA_i_^Me^ binding activities. **(A)** Heat map showing the log2 fold change of translation efficiency (eIF2A^KO^/control cells) of different initiation factors reported to possess binding capacity to tRNA_i_^Met^. Changes in DMSO and Tunicamycin treated samples are shown.

## REFERENCES

Anderson, R., Agarwal, A., Ghosh, A., Guan, B.J., Casteel, J., Dvorina, N., Baldwin, W.M., 3rd, Mazumder, B., Nazarko, T.Y., Merrick, W.C., et al. 2021. eIF2A-knockout mice reveal decreased life span and metabolic syndrome. FASEB journal : official publication of the Federation of American Societies for Experimental Biology 35: e21990. 10.1096/fj.202101105R

Andreev, D.E., O’Connor, P.B., Fahey, C., Kenny, E.M., Terenin, I.M., Dmitriev, S.E., Cormican, P., Morris, D.W., Shatsky, I.N., and Baranov, P.V. 2015. Translation of 5’ leaders is pervasive in genes resistant to eIF2 repression. eLife 4: e03971. 10.7554/eLife.03971

Bohlen, J., Harbrecht, L., Blanco, S., Clemm von Hohenberg, K., Fenzl, K., Kramer, G., Bukau, B., Teleman, A.A. 2020. DENR promotes translation reinitiation via ribosome recycling to drive expression of oncogenes including ATF4. Nature communications 11: 4676. 10.1038/s41467-020-18452-2

Bohlen, J., Roiuk, M., Neff, M., and Teleman, A.A. 2023. PRRC2 proteins impact translation initiation by promoting leaky scanning. Nucleic Acids Res 51: 3391–409. 10.1093/nar/gkad135

Chen, L., He, J., Zhou, J., Xiao, Z., Ding, N., Duan, Y., Li, W., and Sun, L.Q. 2019. EIF2A promotes cell survival during paclitaxel treatment in vitro and in vivo. J Cell Mol Med 23: 6060–71. 10.1111/jcmm.14469

Davies, M.V., and Kaufman, R.J. 1992. The sequence context of the initiation codon in the encephalomyocarditis virus leader modulates efficiency of internal translation initiation. Journal of virology 66: 1924–32. 10.1128/JVI.66.4.1924-1932.1992

Diem, M.D., Chan, C.C., Younis, I., and Dreyfuss, G. 2007. PYM binds the cytoplasmic exon-junction complex and ribosomes to enhance translation of spliced mRNAs. Nature structural & molecular biology 14: 1173–9. 10.1038/nsmb1321

Dmitriev, S.E., Terenin, I.M., Andreev, D.E., Ivanov, P.A., Dunaevsky, J.E., Merrick, W.C., and Shatsky, I.N. 2010. GTP-independent tRNA delivery to the ribosomal P-site by a novel eukaryotic translation factor. The Journal of biological chemistry 285: 26779–87. M110.119693 [pii] 10.1074/jbc.M110.119693

Doench, J.G., Fusi, N., Sullender, M., Hegde, M., Vaimberg, E.W., Donovan, K.F., Smith, I., Tothova, Z., Wilen, C., Orchard, R., et al. 2016. Optimized sgRNA design to maximize activity and minimize off-target effects of CRISPR-Cas9. Nature biotechnology 34: 184–91. 10.1038/nbt.3437

Gaikwad, S., Ghobakhlou, F., Zhang, H., and Hinnebusch, A.G. 2024. Yeast eIF2A has a minimal role in translation initiation and uORF-mediated translational control in vivo. eLife 12. 10.7554/eLife.92916

Golovko, A., Kojukhov, A., Guan, B.J., Morpurgo, B., Merrick, W.C., Mazumder, B., Hatzoglou, M., and Komar, A.A. 2016. The eIF2A knockout mouse. Cell cycle 15: 3115–20. 10.1080/15384101.2016.1237324

Gonzalez-Almela, E., Williams, H., Sanz, M.A., and Carrasco, L. 2018. The Initiation Factors eIF2, eIF2A, eIF2D, eIF4A, and eIF4G Are Not Involved in Translation Driven by Hepatitis C Virus IRES in Human Cells. Front Microbiol 9: 207. 10.3389/fmicb.2018.00207

Grove, D.J., Levine, D.J., and Kearse, M.G. 2023. Increased levels of eIF2A inhibit translation by sequestering 40S ribosomal subunits. Nucleic Acids Res 51: 9983–10000. 10.1093/nar/gkad683

Grove, D.J., Russell, P.J., and Kearse, M.G. 2024. To initiate or not to initiate: A critical assessment of eIF2A, eIF2D, and MCT-1.DENR to deliver initiator tRNA to ribosomes. Wiley interdisciplinary reviews RNA 15: e1833. 10.1002/wrna.1833

Gu, Y., Mao, Y., Jia, L., Dong, L., and Qian, S.B. 2021. Bi-directional ribosome scanning controls the stringency of start codon selection. Nature communications 12: 6604. 10.1038/s41467-021-26923-3

Hinnebusch, A.G., and Lorsch, J.R. 2012. The mechanism of eukaryotic translation initiation: new insights and challenges. Cold Spring Harbor perspectives in biology 4. 10.1101/cshperspect.a011544

Ichihara, K., Matsumoto, A., Nishida, H., Kito, Y., Shimizu, H., Shichino, Y., Iwasaki, S., Imami, K., Ishihama, Y., and Nakayama, K.I. 2021. Combinatorial analysis of translation dynamics reveals eIF2 dependence of translation initiation at near-cognate codons. Nucleic Acids Res 49: 7298–317. 10.1093/nar/gkab549

Jaafar, Z.A., Oguro, A., Nakamura, Y., and Kieft, J.S. 2016. Translation initiation by the hepatitis C virus IRES requires eIF1A and ribosomal complex remodeling. eLife 5. 10.7554/eLife.21198

Kim, E., Kim, J.H., Seo, K., Hong, K.Y., An, S.W.A., Kwon, J., Lee, S.V., and Jang, S.K. 2018. eIF2A, an initiator tRNA carrier refractory to eIF2alpha kinases, functions synergistically with eIF5B. Cell Mol Life Sci 75: 4287–300. 10.1007/s00018-018-2870-4

Kim, J.H., Park, S.M., Park, J.H., Keum, S.J., and Jang, S.K. 2011. eIF2A mediates translation of hepatitis C viral mRNA under stress conditions. The EMBO journal 30: 2454–64. 10.1038/emboj.2011.146

Komar, A.A., Gross, S.R., Barth-Baus, D., Strachan, R., Hensold, J.O., Goss Kinzy, T., and Merrick, W.C. 2005. Novel characteristics of the biological properties of the yeast Saccharomyces cerevisiae eukaryotic initiation factor 2A. The Journal of biological chemistry 280: 15601–11. 10.1074/jbc.M413728200

Komar, A.A., and Merrick, W.C. 2020. A Retrospective on eIF2A-and Not the Alpha Subunit of eIF2. International journal of molecular sciences 21. 10.3390/ijms21062054

Kozak, M. 1991. Structural features in eukaryotic mRNAs that modulate the initiation of translation. The Journal of biological chemistry 266: 19867–70.

Kwon, O.S., An, S., Kim, E., Yu, J., Hong, K.Y., Lee, J.S., and Jang, S.K. 2017. An mRNA-specific tRNAi carrier eIF2A plays a pivotal role in cell proliferation under stress conditions: stress-resistant translation of c-Src mRNA is mediated by eIF2A. Nucleic Acids Res 45: 296–310. 10.1093/nar/gkw1117

Labun, K., Montague, T.G., Krause, M., Torres Cleuren, Y.N., Tjeldnes, H., and Valen, E. 2019. CHOPCHOP v3: expanding the CRISPR web toolbox beyond genome editing. Nucleic Acids Res 47: W171–W74. 10.1093/nar/gkz365

Levin, D.H., Kyner, D., and Acs, G. 1973. Protein initiation in eukaryotes: formation and function of a ternary complex composed of a partially purified ribosomal factor, methionyl transfer RNA, and guanosine triphosphate. Proceedings of the National Academy of Sciences of the United States of America 70: 41–5. 10.1073/pnas.70.1.41

Liang, H., He, S., Yang, J., Jia, X., Wang, P., Chen, X., Zhang, Z., Zou, X., McNutt, M.A., Shen, W.H., et al. 2014. PTENalpha, a PTEN isoform translated through alternative initiation, regulates mitochondrial function and energy metabolism. Cell metabolism 19: 836–48. 10.1016/j.cmet.2014.03.023

Lowe, D.D., and Montell, D.J. 2022. Unconventional translation initiation factor EIF2A is required for Drosophila spermatogenesis. Developmental dynamics : an official publication of the American Association of Anatomists 251: 377–89. 10.1002/dvdy.403

Merrick, W.C., and Anderson, W.F. 1975. Purification and characterization of homogeneous protein synthesis initiation factor M1 from rabbit reticulocytes. The Journal of biological chemistry 250: 1197–206.

Oertlin, C., Lorent, J., Murie, C., Furic, L., Topisirovic, I., and Larsson, O. 2019. Generally applicable transcriptome-wide analysis of translation using anota2seq. Nucleic Acids Res 47: e70 10.1093/nar/gkz223

Palmiter, R.D. 1975. Quantitation of parameters that determine the rate of ovalbumin synthesis. Cell 4: 189. 10.1016/0092-8674(75)90167-1

Panzhinskiy, E., Skovsø, S., Cen, H.H., Chu, K.Y., MacDonald, K., Soukhatcheva, G., Dionne, D.A., Hallmaier-Wacker, L.K., Wildi, J.S., Marcil, S., et al. 2021. Eukaryotic translation initiation factor 2A protects pancreatic beta cells during endoplasmic reticulum stress while rescuing translation inhibition. bioRxiv: 2021.02.17.431676. 10.1101/2021.02.17.431676

Pestova, T.V., and Kolupaeva, V.G. 2002. The roles of individual eukaryotic translation initiation factors in ribosomal scanning and initiation codon selection. Genes Dev 16: 2906–22. 10.1101/gad.1020902

Reineke, L.C., Komar, A.A., Caprara, M.G., and Merrick, W.C. 2008. A small stem loop element directs internal initiation of the URE2 internal ribosome entry site in Saccharomyces cerevisiae. The Journal of biological chemistry 283: 19011–25. 10.1074/jbc.M803109200

Reineke, L.C., and Merrick, W.C. 2009. Characterization of the functional role of nucleotides within the URE2 IRES element and the requirements for eIF2A-mediated repression. Rna 15: 2264–77. 10.1261/rna.1722809

Roiuk, M., Neff, M., and Teleman, A.A. 2024. eIF4E-independent translation is largely eIF3d-dependent. Nature communications 15: 6692. 10.1038/s41467-024-51027-z

Sanz, M.A., Almela, E.G., Garcia-Moreno, M., Marina, A.I., and Carrasco, L. 2019. A viral RNA motif involved in signaling the initiation of translation on non-AUG codons. Rna 25: 431–52. 10.1261/rna.068858.118

Sanz, M.A., Gonzalez Almela, E., and Carrasco, L. 2017. Translation of Sindbis Subgenomic mRNA is Independent of eIF2, eIF2A and eIF2D. Scientific reports 7: 43876. 10.1038/srep43876

Schleich, S., Acevedo, J.M., Clemm von Hohenberg, K., and Teleman, A.A. 2017. Identification of transcripts with short stuORFs as targets for DENR*MCTS1-dependent translation in human cells. Scientific reports 7: 3722. 10.1038/s41598-017-03949-6

Schleich, S., Strassburger, K., Janiesch, P.C., Koledachkina, T., Miller, K.K., Haneke, K., Cheng, Y.S., Kuchler, K., Stoecklin, G., Duncan, K.E., et al. 2014. DENR-MCT-1 promotes translation re-initiation downstream of uORFs to control tissue growth. Nature 512: 208–12. 10.1038/nature13401

Sendoel, A., Dunn, J.G., Rodriguez, E.H., Naik, S., Gomez, N.C., Hurwitz, B., Levorse, J., Dill, B.D., Schramek, D., Molina, H., et al. 2017. Translation from unconventional 5’ start sites drives tumour initiation. Nature 541: 494–99. 10.1038/nature21036

Shafritz, D.A., and Anderson, W.F. 1970. Isolation and partial characterization of reticulocyte factors M1 and M2. The Journal of biological chemistry 245: 5553–9.

Sidrauski, C., McGeachy, A.M., Ingolia, N.T., and Walter, P. 2015. The small molecule ISRIB reverses the effects of eIF2alpha phosphorylation on translation and stress granule assembly. eLife 4. 10.7554/eLife.05033

Sonobe, Y., Ghadge, G., Masaki, K., Sendoel, A., Fuchs, E., and Roos, R.P. 2018. Translation of dipeptide repeat proteins from the C9ORF72 expanded repeat is associated with cellular stress. Neurobiology of disease 116: 155–65. 10.1016/j.nbd.2018.05.009

Starck, S.R., Jiang, V., Pavon-Eternod, M., Prasad, S., McCarthy, B., Pan, T., and Shastri, N. 2012. Leucine-tRNA initiates at CUG start codons for protein synthesis and presentation by MHC class I. Science 336: 1719–23. 10.1126/science.1220270

Starck, S.R., Tsai, J.C., Chen, K., Shodiya, M., Wang, L., Yahiro, K., Martins-Green, M., Shastri, N., and Walter, P. 2016. Translation from the 5’ untranslated region shapes the integrated stress response. Science 351: aad3867. 10.1126/science.aad3867

Tahmasebi, S., Khoutorsky, A., Mathews, M.B., and Sonenberg, N. 2018. Translation deregulation in human disease. Nature reviews Molecular cell biology 19: 791–807. 10.1038/s41580-018-0034-x

Tusi, S.K., Nguyen, L., Thangaraju, K., Li, J., Cleary, J.D., Zu, T., and Ranum, L.P.W. 2021. The alternative initiation factor eIF2A plays key role in RAN translation of myotonic dystrophy type 2 CCUG*CAGG repeats. Human molecular genetics 30: 1020–29. 10.1093/hmg/ddab098

Zoll, W.L., Horton, L.E., Komar, A.A., Hensold, J.O., and Merrick, W.C. 2002. Characterization of mammalian eIF2A and identification of the yeast homolog. The Journal of biological chemistry 277: 37079–87. 10.1074/jbc.M207109200

